# Acetate supplementation rescues social deficits and alters transcriptional regulation in prefrontal cortex of Shank3 deficient mice

**DOI:** 10.1101/2020.04.29.065821

**Authors:** Aya Osman, Nicholas L. Mervosh, Ana N. Strat, Tanner J. Euston, Gillian Zipursky, Rebecca M. Pollak, Katherine R. Meckel, Scott R. Tyler, Kenny L. Chan, Ariela Buxbaum Grice, Elodie Drapeau, Lev Litichevskiy, Jasleen Gill, Christoph A. Thaiss, Joseph. D. Buxbaum, Michael S. Breen, Drew D. Kiraly

## Abstract

Autism spectrum disorder (ASD) is a heterogenous neurodevelopmental disorder with complex pathophysiology including both genetic and environmental factors. Recent evidence demonstrates the gut microbiome and its resultant metabolome can influence brain and behavior and have been implicated in ASD. To investigate gene by microbiome interactions in a model for genetic risk of ASD, we utilize mutant mice carrying a deletion of the ASD-associated *Shank3* gene (Shank3^KO^). *Shank3*^KO^ have altered microbiome composition and function at baseline in addition to social deficits. Further depletion of the microbiome with antibiotics exacerbates social deficits in *Shank3*^KO^, and results in transcriptional changes in the frontal cortex. Supplementation with the microbial metabolite acetate leads to reversal of social behavioral phenotypes even in mice with a depleted microbiome, and significantly alters transcriptional regulation in the prefrontal cortex. These results suggest a key role for the gut microbiome and the neuroactive metabolite acetate in regulating ASD-like behaviors.

## Introduction

Autism spectrum disorder (ASD) is a potentially serious neurodevelopmental disorder characterized by deficits in social function and stereotyped and repetitive behaviors^1^. The prevalence of ASD is remarkably high, with recent estimates approximating 1 in 44 children aged 8 and under affected^2^. Despite this, there are currently no disease modifying pharmacotherapies for treatment of ASD. This is in part due to a complex pathophysiology that is driven by combination of genetic and environmental risk factors. Large scale genetic studies have identified more than 100 gene loci associated with increased risk of ASD^3^, and a number of single gene causes of ASD have been identified (e.g. *Fmr1*, *Shank3*)^4,5^. While host genetics is a key factor in ASD, a more comprehensive understanding of non-genetic environmental contributions and how to mitigate them represents an attractive area of research to understand and reduce the development and progression of ASD.

A growing body of evidence suggests that the resident bacteria of the gastrointestinal tract, collectively known as the gut microbiome, play a key role in host brain development and behavior through a mechanism known as the gut-brain axis^6^, which has been implicated in neurodevelopmental disorders. Disruptions to the gut microbiome lead to alterations in gene expression and epigenetic regulation in the brain as well as alterations in neuronal activity patterns and synaptic plasticity in multiple limbic substructures^7–9^. Clinical data examining patients with ASD compared to neurotypical controls has corroborated changes in microbiome composition^10–12^ and metabolic profile^13,14^. However, specific microbial and metabolic changes associated with a diagnosis have varied between studies. Additionally, patients with both idiopathic and syndromic forms of ASD report gastrointestinal issues at rates much higher than the general population^5,12^.

Evidence linking the microbiome to ASD is further supported by findings from murine models for ASD. Early studies showed that germ-free mice (i.e. mice that are raised completely sterile with no internal or external microbiome) have marked deficits in social behaviors and in gene expression profiles in multiple brain regions related to regulation of these behaviors^15–17^. Additionally, mice who have had their microbiomes depleted with antibiotics show deficits in social behavior, cognitive flexibility, and multiple other ASD-like behaviors^7,18,19^. At a cellular and molecular level, disruption of the microbiome with prolonged antibiotics leads to changes in neuronal and glial structure, changes in cell firing patterns, and altered transcriptional and epigenetic signatures in multiple brain structures^7,20–22^.

Although the exact mechanisms by which gut bacteria influence the brain are not completely understood, the strongest literature suggests the production of neuroactive metabolites by the microbiome as a key gut-brain signaling mechanism^7,22,23^. Among the best studied metabolites in this gut-brain axis are the short-chain fatty acids (SCFA) which are produced in the process of bacterial fermentation of fiber and can cross the blood-brain barrier and influence chromatin structure and gene expression among other effects^24,25^.

While the literature demonstrating a gut-brain connection in ASD is robust and continues to grow, there are few studies specifically modeling gene by microbiome interactions using available mouse models. An attractive gene candidate for this line of research is the *Shank3* gene which encodes a synaptic scaffolding protein and is the causative gene underlying Phelan McDermid Syndrome (PMS). Importantly, patients with PMS manifest features of ASD with cognitive impairment, and also have very high rates of gastrointestinal issues^5^ – suggesting a potential for a gut brain link. Indeed, several studies have examined the microbiota in mouse lines lacking specific *Shank3* isoforms and have found changes in the gut microbiome composition that relate to ASD-like phenotypes^26,27^.

Here we utilize a recently characterized mouse model with a deletion of exons 4-22 of the *Shank3* gene resulting in deletion of all functional isoforms of *Shank3* (*Shank3^KO^*)^28^. Using 16S and metagenomic sequencing, and targeted metabolomics we identify changes in microbiome form and function driven by *Shank3* genotype. Additionally, using antibiotic induced microbiome depletion we see an exacerbation in metabolic alterations and that *Shank3^KO^* mice are particularly sensitive to behavioral effects of microbiome disruption. Using RNA-sequencing of the frontal cortex we identify a unique gene expression signature induced by microbiome depletion in the *Shank3^KO^* mice. Finally, targeting and replenishing depleted levels of the SCFA acetate were found to reverse social deficits and markedly altered cortical transcriptional regulation even in mice lacking a complex microbiome. Taken together, our data build on existing work identifying important gene by microbiome interactions in ASD and provide a preliminary mechanism of this interaction via identification of a specific metabolite.

## Materials and Methods

### Mice

All animal procedures were approved by the Institutional Animal Care and Use Committee of the Icahn School of Medicine at Mount Sinai. *Shank3^KO^* mice who are homozygous for the Shank3 gene were generated from breeders heterozygous *Shank3* gene^2^, and littermate mice were used for all experiments. Both male and female mice were weaned at 21 days of age and caged according to sex with at least one littermate from each genotype, with cages containing 2-5 littermates per cage. Food was available ad libitum throughout the experiments, and drinking water was modified as described below. The mouse holding room was maintained at a constant temperature of 21°C and 55% Humidity with a 12-hour light dark cycle (on 0700 / off 1900).

### Drink Solutions

Immediately after weaning, mice were split equally into control or Abx treatment groups for Experiment 1 (**Fig. 2**) and into control, Abx, Acetate, or Abx + acetate (Ab/Ac) treatment groups for experiment 2 (**Fig. 4**). Control mice were provided with standard drinking water from the tap. Abx mice received drinking water containing a non-absorbable antibiotic cocktail (Abx - Neomycin 2 mg/ml, Pimaricin 1.2 ug/ml, Vancomycin 0.2 g/ml, and Bacitracin 0.5 mg/ml)^22^. Acetate treatment groups received drinking water containing 100 mM acetate (Sigma #S5636). Abx+Ac groups received Abx and Acetate at the described concentrations. Mice were maintained on these treatments from PND21 till the end of behavioral testing on approximately PND65. Bottle as well as mouse weights were recorded biweekly for first two weeks of treatment for a subset of mice, and then weekly afterwards to ensure mice were maintaining health. Antibiotic containing solutions were changed weekly to maintain antibiotic efficacy.

### 16s-Sequencing

Bacterial genomic DNA was isolated from frozen cecal samples using the QiaAmp DNA stool kit (Qiagen) according to standard protocols with a modification that included an additional step of bead beating to ensure uniform lysis of bacterial cells^63^. Extracted DNAs were checked for quality and quantity by spectrophotometric measurements with NanoDrop (ThermoFisher Scientific Inc) and stored at −20C until processed for amplification. PCR amplification used primers 341F (5’-CCTACGGGNGGCWGCAG-3’) and 805R (5’-GACTACHVGGGTATCTAATCC-3’) targeting regions (V3-V4) of the 16S rRNA gene. The libraries were sequenced using illumina NovaSeq (2 × 250 bp paired-end) platform using standard parameters.

#### 16S Sequencing Data Analysis

Amplicons were trimmed, merged using FLASH^10^ and chimera filtered using Vsearch (v2.3.4). Sequences with ≥97% similarity were assigned to the same operational taxonomic units (OTUs). Representative sequences were chosen for each OTU, followed by taxonomic assignment using the RDP (Ribosomal Database Project) classifier^63^. The differences of the dominant species in different groups and multiple sequence alignment were conducted by mafft software (v7.310). Alpha diversity for analyzing complexity of species diversity were calculated by Quantitative Insights Into Microbial Ecology (QIIME) v1.8.0^64^. Principle coordinates analysis plots were generated using the Unifrac distance as an assessment of beta diversity, also using the (QIIME) package. Statistical significance for alpha diversity was analysed using a 2×2 between subjects ANOVA with genotype and drink solution provided as fixed factors.

### Shotgun Metagenomic Sequencing and Data Analysis

Bacterial genomic DNA was isolated from frozen cecal samples using the QiaAmp DNA stool kit (Qiagen) as described above was subjected to metagenomic sequencing. Metagenomic data were pre-processed using Sunbeam^65^. In brief, reads were trimmed of adapters using cutadapt. Reads were quality-trimmed using trimmomatic^66^. Low-complexity reads were removed using Komplexity [https://github.com/eclarke/komplexity], and reads aligning to the mouse genome were removed using BWA^67^. DIAMOND^68^ was used to align reads to all prokaryotic protein sequences from KEGG^69^. KEGG Orthologs (KOs) that received fewer than 30 counts across all samples were removed. KEGG pathway counts were computed by summing the counts for all KOs belonging to each pathway. KOs and pathways with differential abundance between WT/HET and KO samples were identified using DESeq2^70^. A significance threshold of FDR *p* < 0.2 was applied. Absolute counts were provided to DESeq2, but relative counts (each count divided by the sum of per-sample counts) were used for box plots.

### Cecal Metabolomics

Cecal contents were collected and flash frozen on dry ice and then stored at −80°C until processing. Samples were processed by the metabolomics core at University of Pennsylvania according to standard protocols. Short chain fatty acids and amino acids were quantified using a Water Acquity uPLC System with a Photodiode Array Detector and an autosampler (192 sample capacity). For short chain fatty acids samples were analyzed on a HSS T3 1.8 μm 2.1×150 mm column with a flow rate of 0.25 mL/min, an injection volume of 5 uL, a column temperature of 40°C, a sample temperature of 4°C, and a run time of 25 min per sample. Eluent A was 100 mM sodium phosphate monobasic, pH 2.5; eluent B was methanol; the weak needle wash 0.1% formic acid in water; the strong needle wash 0.1% formic acid in acetonitrile, and the seal wash is 10% acetonitrile in water. The gradient was 100% eluent A for 5 min, gradient to 70% eluent B from 5-22 min, and then 100% eluent A for 3 min. The photodiode array is set to read absorbance at 215 nm with 4.8 nm resolution. Amino acid concentrations were quantified via ultra-performance liquid chromatography (Waters Acquity UPLC system) with an AccQ-Tag Ultra C18 1.7μm 2.1×100mm column and a photodiode detector array. Analysis was performed using the UPLC AAA H-Class Application Kit (Waters Corporation, Milford, MA) according to manufacturer’s instructions.

For both assays, standards were run at the beginning and end of each metabolomics run. Quality control checks (blanks and standards) were run every eight samples. Results were rejected if the standards deviate by greater than ±5%. Samples were quantified against standard curves of at least five points run in triplicate. All chemicals and reagents are mass spectrometry grade.

### Behavioral assessments

Behavioral experiments were conducted between 10:00 am and 18:00 during the light phase of the light/dark cycle in soundproof cubicles designated for behavioral testing under red lights. Mice were brought to the room of the testing area an hour prior to testing in order to habituate to the new environment. Mice from genotype and treatment groups were tested on the same day in random order to control for any potential circadian influences.

#### Open field

Mice were tested in a square arena (45 x 45 cm) and enclosed by 25cm high walls (Plexiglass). The test mouse was placed in center of arena and allowed to move freely for 10 minutes or 1 hour. Mice activity was recorded by video tracking (Noldus Ethovision). The arena was virtually divided into central and peripheral regions using the Noldus software. The total distance travelled, as well as the total time spent in the center were calculated.

#### Elevated Zero Maze

Mice were tested in an annular black Plexiglas runway, 5 cm wide, 60 cm in diameter and raised 60 cm off the ground (Maze Engineers). The runway was divided equally into four equal quadrants. Two opposite quadrants were “open” enclosed only by a 1 cm inch lip; the remaining two quadrants were “closed” surrounded by a 25 cm high dark, opaque walls. Outer walls were constructed of dark grey plastic, inner walls were black Plexiglas. All test subject was placed in the centre of an open arm and allowed to move freely for 5 minutes. Mouse activity was recorded by video tracking (Noldus Ethovision). The arena was virtually divided into open and closed regions using the Noldus software. The duration frequency of entries into the open arms were calculated.

#### Three chambered Social Interaction Test

Sociability was tested in a three chambered apparatus according to standard procedures^71^. The test consisted of three stages. The first two were for habituation to the chamber, and the third was the measured test phase. During the first stage the test subject was placed into the central neutral chamber with both dividers closed and was allowed to explore the middle chamber for 5 minutes. During the second stage of the test, the two dividers were removed, and the test subject explored all three chambers for 10 minutes without another mouse present. During the final stage of the test the mouse was returned to the middle chamber with both dividers closed, and an empty wire cup and a wire cup containing a novel interactor mouse were placed in the two outside chambers. The wire cups allow for olfactory, visual, auditory and tactile contact contacts between mice but not sexual contact or fighting. The doors between chambers were then opened, and the test subject was allowed to explore all three chambers again for 10 minutes. All three sessions were video tracked (Noldus Ethovision) and the amount of time spent in each chamber as well as the time spent interacting with either empty cup or unfamiliar mouse was automatically calculated. If a mouse spent 70% more time in one chamber than the other during the habituation stage (stage 2) they were dropped from the analysis. Mice used as interactors were adult C57BL/6J mice of the same age and sex as the test mouse. Interactor mice were ordered from Jackson labs and allowed at least one week to habituate to the colony room prior to any testing. Twenty-four hours to testing, the interactor mice were habituated to the interaction apparatus and the wire cup to prevent any confounding effects of novelty stress.

#### Behavior Statistical analysis

All behavioral analyses were performed using GraphPad Prism with two-way ANOVAs with Holm-Sidak post hoc tests as appropriate for 2 x 2 experiments, and as two-tailed T-tests for pairwise comparisons.

### RNA-Sequencing

Medial Prefrontal cortex (mPFC) samples were rapidly dissected from ~60-day old mice and flash frozen on dry ice. Total RNA was isolated by homogenizing tissue in Qiazol (Qiagen) using a motorized pellet pestle and purified using RNeasy mini kits from Qiagen (74106) with optional on-column DNAase digestion (79254) per manufacturer’s protocol. The integrity and purity of total RNA were assessed using Agilent Bioanalyzer and OD260/280 using Nanodrop. The RNA sequencing library was generated using NEBNext Ultra II Directional RNALibrary Prep Kit for Illumina using manufacturer’s instructions (New England Biolabs, Ipswich, MS, USA). The sequencing library was validated on the Agilent TapeStation (Agilent Technologies, Palo Alto, CA, USA) and quantified by using Qubit 2.0 Fluorometer (ThermoFisher Scientific, Waltham, MA, USA) as well as by quantitative PCR (KAPA Biosystems, Wilmington, MA, USA). The libraries were then multiplexed and sequenced on an Illumina HiSeq 4000 instrument using a 2×150bp paired-end configuration according to manufacturer’s instructions. Image analysis and base calling were conducted by the HiSeq Control Software (HCS). Raw sequence data (.bcl files) generated from Illumina HiSeq were converted into fastq files and de-multiplexed using Illumina bcl2fastq 2.20 software. One mismatch was allowed for index sequence identification.

#### RNA-sequencing Data Analysis

The raw RNA-Seq reads for each sample were aligned and read counts generated using the cloud-based BioJupies software package using the default settings^72^. Raw count data measured at least 32,544 genes across all samples. Nonspecific filtering required more than 1 count per million (cpm) in at least 3 samples and retained 14,870 genes. Filtered raw count data was then subjected to voom normalization^73^. Normalized data were inspected for outlying samples using unsupervised clustering of samples (Pearson’s correlation coefficient and average distance metric) and principal component analysis to identify outliers outside two standard deviations from these grand averages. Based on these metrics, no significant outliers were found. Differential gene expression (DGE) signatures between samples were identified using moderated *t*-test in the limma package^73^. The following covariates were included in the model to adjust for their potential confounding influence on gene expression between group main effects: treatment group, sex and genotype. A surrogate variable analysis was also conducted to remove any hidden variables besides treatment, sex and genotype on DGE signatures^3^. Statistical significance threshold was set as a nominal *p*-value of < 0.01 to permit sufficient enough genes to move forward with functional characterization. Identification of significantly enriched gene ontologies and upstream regulators was performed using G:Profiler and Ingenuity Pathway Analysis (IPA) software packages as indicated^74,75^.

### Mouse Serum Metabolomics

Serum was isolated from trunk blood and stored at −80°C until processing. Targeted short chain fatty acid and untargeted metabolomic analysis were performed using liquid chromatography-mass spectrometry (LC/MS). Chemical isotope labelling of each sample was carried out by following the SOPs provided in the labeling kits (Nova Medical Testing Inc., Product Numbers: NMT-4101-KT, NMT-4123-KT, NMT-4145-KT, and NMT-4167-KT) to measure the total concentration of metabolites in a 25 μL aliquot of sample^76^. Based on the total concentration of labelled metabolites, each sample was then normalized to the same concentration of 1.2mM^77^. An Agilent eclipse plus reversed-phase C18 column (150 × 2.1 mm, 1.8 μm particle size) was then used to separate the labelled samples and Agilent 1290 LC linked to Bruker Impact II QTOF Mass Spectrometer was used to detect the labelled metabolites. Mobile phase A was 0.1% (v/v) formic acid in water and mobile phase B was 0.1% (v/v) formic acid in acetonitrile. The LC gradient setting was: t = 0 min, 25% B; t = 10 min, 99% B; t = 15 min, 99% B; t = 15.1 min, 25% B; t = 18 min, 25% B. The flow rate was 400 μL/min. The column oven temperature was 40 °C, mass spectral acquisition rate was 220 to 1000 m/z. Raw mass data were exported to csv file using DataAnalysis 4.4 (Bruker Daltonics). Data analysis was performed using IsoMS Pro 1.2.12 (Nova Medical Testing Inc.)^78^. Metabolite identification was subsequently carried out using a two-tiered approach against NovaMT Metabolite Databases 2.0 (Nova Medical Testing Inc.). In tier 1 metabolites were searched against the in-house mass spec labelled metabolite library based on accurate mass and retention time. In tier 2, the remaining peak pairs were searched against a linked identity library based on accurate mass and predicted retention time matched^79^.

#### Metabolomics Data Analysis

For metabolomic data analysis, all data was firstly normalized and re-scaled to set the median equal to one. Next, any missing values were imputed to the minimum scaled value for each metabolite. Statistical analyses were performed in Excel and other software. Principle Component Analysis (PCA) was performed with (MetaboAnalyst 5.0^80^. The heat map was generated using the Log Fold change of metabolites identified in tier 1 and tier 2 analysis. For univariate analysis, a student’s t-test was used to calculate significance of metabolites (*p*<0.05). FDR adjusted p-values were calculated using Store’s q-value (q < 0.25).

### Human Plasma Metabolomics

We leveraged our published plasma metabolomic profiles derived from 32 PMS probands and 28 controls (siblings *n*=27; parent *n*=1) ascertained at the Seaver Autism Center for Research and Treatment at the Icahn School of Medicine at Mount Sinai^4^. For these samples, a targeted panel of eight short chain fatty acid metabolites were generated. Data quality control was conducted as previously described^4^. In brief, any missing values were imputed to the minimum scaled value for each metabolite. Data were normalized by log_2_ transformation prior to performing any statistical analysis.

### Human metabolomics statistical analyses

A moderated t-test from limma was used to assess the effect of categorical variables such as genotype, variant class, variant type, medication status, ADOS criteria, ADI criteria and GI status on each of the eight SCFAs in PMS patient compared to controls. Correlative analysis was also conducted using limma to assess the effect of quantitative variables: log deletion size, raw deletion size, number of medications, IQ/DQ, ADOS variables, RRB score, ADI variables, Vineland variables and ABC variables. Empirical Bays method was used to calculate moderated t and F statistics for both types of analysis. These analyses were adjusted for possible influence of demographic variables, age and race. Adjusted *p* <0.05 was accepted as statistically significant.

### Figures

Graphs of all figures were created in Graphpad Prism and R. Experimental timelines were generated in BioRender with full permission to publish.

## Results

### Deletion of the Shank3 Gene Results in a Dysregulated Gut Microbiome and Metabolic Output

To identify effects of *Shank3* gene deletion on microbiome composition and metabolic output, both 16S ribosomal and shotgun metagenomic sequencing was performed on cecal contents of untreated Wt and *Shank3^KO^* mice in adulthood. To control for potential maternal and environmental confounds, analysis was performed on littermate mice co-housed by genotype. Measures of alpha diversity, which calculates the richness and evenness of species in a microbial sample^29^, show decreased alpha diversity in *Shank3^KO^* mice when calculated by both the Simpson (**Fig. 1A** – *p* = 0.01) and Shannon (**Fig. 1B** – *p* = 0.03) diversity indices. This decrease in diversity is apparent when the relative expression of bacterial phyla in the microbiomes of the two genotypes are qualitatively compared (**Fig. 1C**). In mammals the most abundant bacterial phyla in the gut microbiome are the Bacteroidetes and Firmicutes, and their presence and relative abundance have been tied to the health of the microbiome^30^. *Shank3^KO^* mice have a significant decrease in the abundance of bacteria from the Firmicutes phylum (**Fig. 1D** – *p* = 0.003), and a trend toward increase in bacteria from the Bacteroidetes phylum (**Fig. 1E** – *p* = 0.08). When the ratio of these species from each mouse is compared, *Shank3^KO^* mice have a significant decrease in the Firmicutes/Bacteroidetes ratio (**Fig. 1F** – *p* = 0.02).

**Figure 1:**
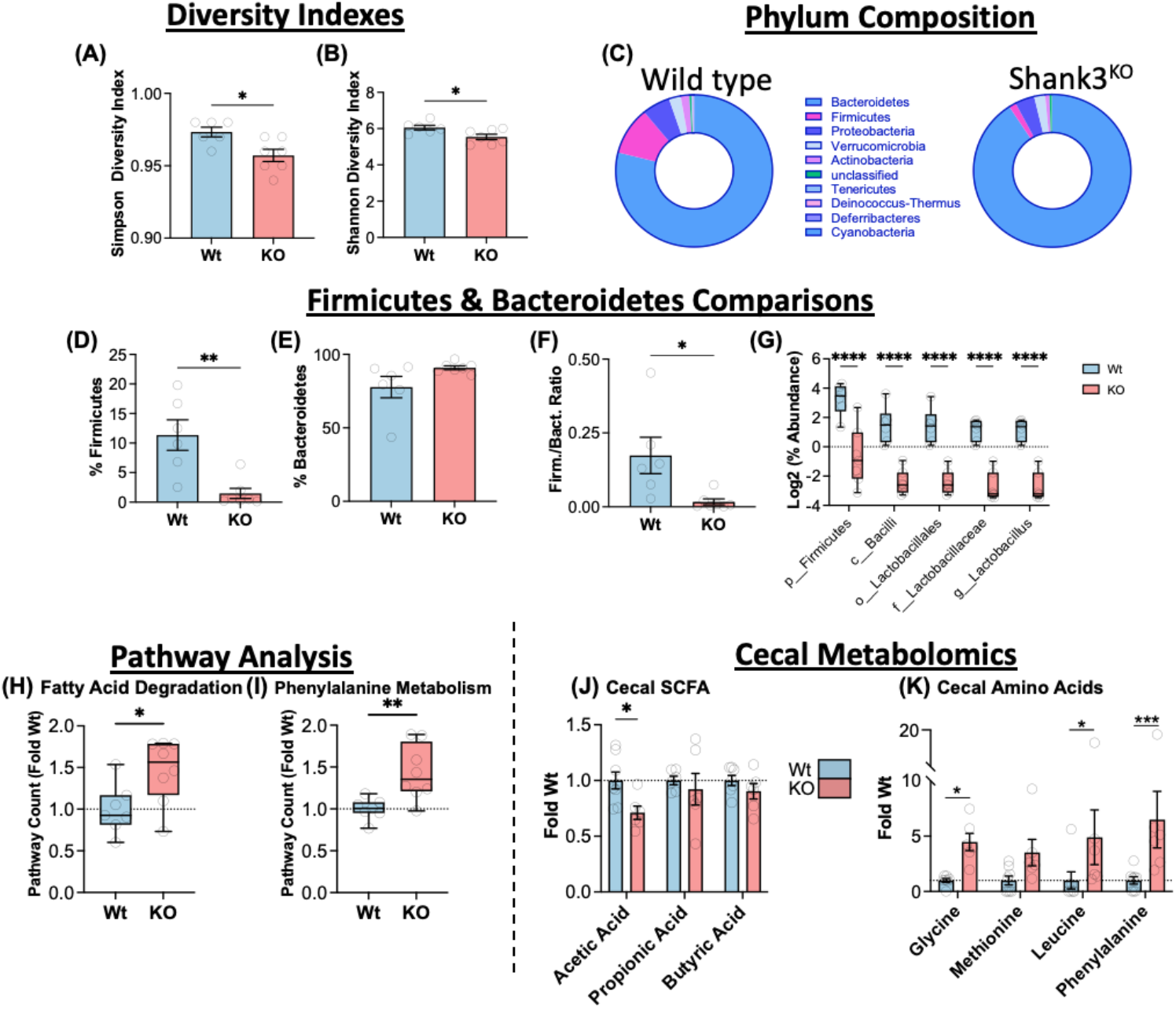
Deletion of the Shank3 Gene Results in a Dysregulated Gut Microbiome and Metabolome. Shank3 ^KO^ mice show decreased microbiome diversity by Simpson **(A)** and Shannon **(B)** diversity metrics. **(C)** Donut charts showing the relative Phylum composition of Wt and Shank3^KO^ mice. **(D)** Relative composition of bacteria from the Firmicutes phylum was decreased in Shank3^KO^, **(E)** but no significant difference in Bacteroidetes. **(F)** The ratio of Firmicutes to Bacteroidetes is significantly decreased in Shank3^KO^ mice. **(G)** Relative abundance of phylogenies containing *Lactobacillus* showed decreases across all taxonomic levels. Functional pathway counts for Fatty Acid degradation **(H)** and Phenylalanine metabolism **(I)** were both significantly increased in Shank3^KO^ mice. (**J**) Targeted metabolomics shows levels of Short Chain Fatty Acid (SCFA) Acetic Acid are significantly decreased in Shank3^KO^ mice. **(K)** Levels of amino acids Glycine, Phenylalanine, and Leucine were increased in Shank3^KO^ mice. All data presented as mean ± SEM \**p*<0.05 \*\**p*<0.01 \*\*\**p*<0.001. *n* = 6-8 mice per group. Both male and female mice used in equal ratios.

**Figure 2:**
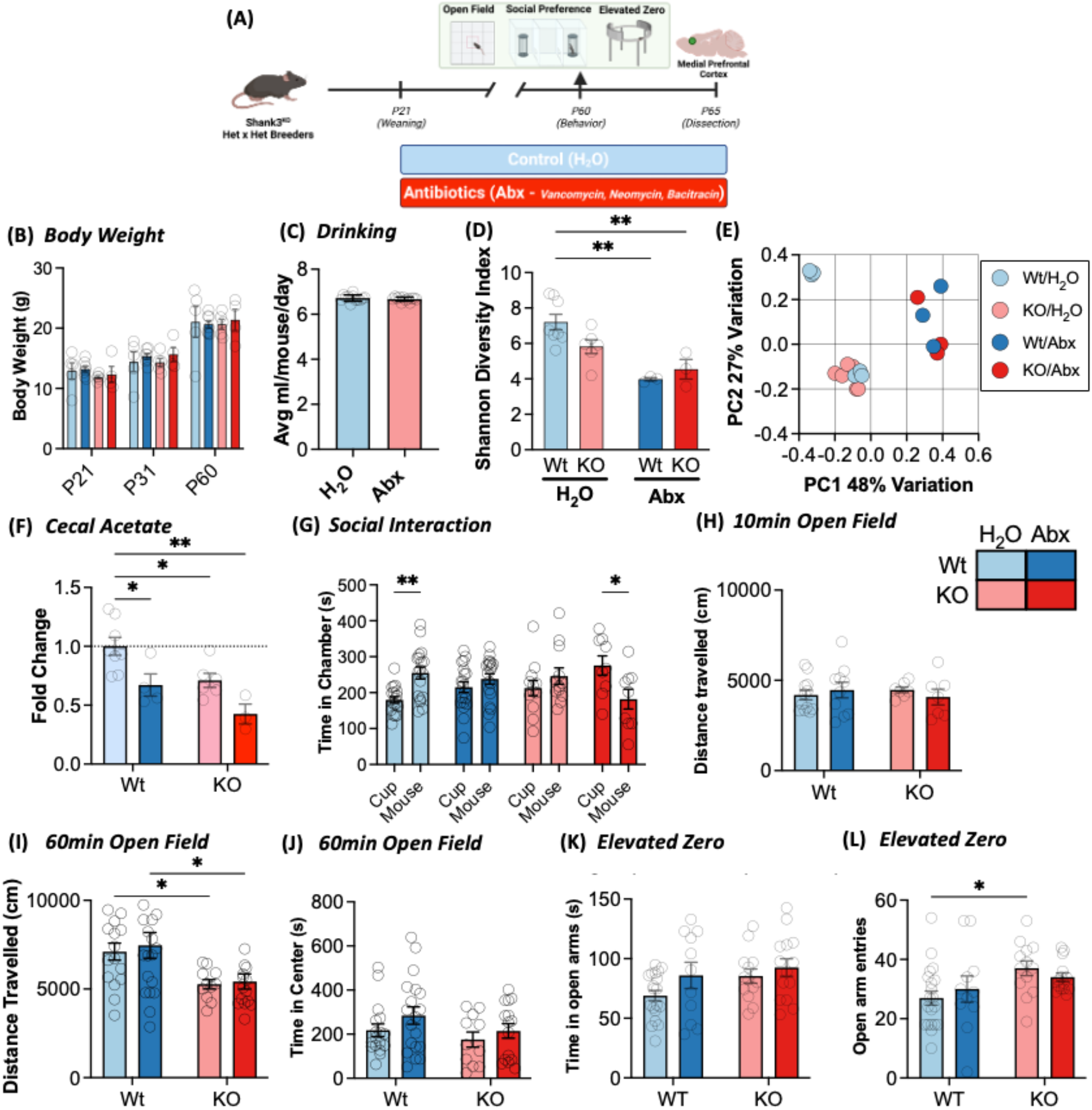
Antibiotic Treatment Throughout Development Exacerbates Metabolic and Social Deficits Caused by Shank3 Deletion. **(A)** Study design for Experiment 1 and mPFC collection. **(B)** Wild-type and Shank3^KO^ mice show normal body weight growth over time regardless of genotype or treatment and **(C)** drink the same amount of control or Abx treated water. **(D)** Antibiotics results in marked depletion of microbial diversity in both Wt and Shank3^KO^ mice. **(E)** Unweighted PCOA plot of beta diversity shows a marked shift induced by antibiotic treatment in both genotypes. **(F)** Levels of Acetic Acid were decreased in control Shank3^KO^ mice and further decreased by Abx treatment. **(G)** In the three-chambered social interaction task, Wt-H_2_O mice spent increased time in the chamber containing the novel social interactor while Shank3 ^KO^-H_2_O did not show a significant preference for the social interactor. Treatment with Abx caused Wt mice to lose social preference and exacerbated social deficits in KO mice. **(H)** Wt and Shank3^KO^ mice of both treatment groups exhibit similar amounts of locomotor activity during a ten-minute open field task. **(I)** Shank3^KO^ mice show decreased locomotion in 60-minute open field not affected by antibiotics. **(J)** No genotype or treatment differences are seen on time in center of an open field. **(K)** In the elevated O-maze no significant differences in total time spent in the open arms in an elevated zero task. **(L)** Shank3^KO^-H_2_O mice display increased frequency of entries into the open arm of the elevated O-maze. All data presented as mean ± SEM. \**p* < 0.05; \*\**p* < 0.01; \*\*\**p* < 0.001; \*\*\*\**p* < 0.0001]. *n* = 3-19

To refine understanding of *Shank3* genotype effects on microbiome composition, examination of phylogeny within Firmicutes down to the species level was conducted. Members of the class Bacilli, order Lactobacillales, family *Lactobacillaceae*, and genus *Lactobacillus* were significantly reduced in *Shank3^KO^* mice (**Fig. 1G** - main effect of genotype F_(1,55)_=196.4 *p* < 0.001 and main effect of taxonomical class F_(4,55)_=7.990; *p* < 0.001; all significant by post-hoc testing *p* < 0.001). A list of all bacterial changes assessed at each taxonomical level is available as **Supplemental Table 1.** Predicted functional consequences of these changes were analyzed with shotgun metagenomics followed by KEGG pathway analysis. From these analyses there were 37 pathways that met statistical significance (*p adj*.<0.2). Of these, Fatty Acid (FA) degradation (**Fig. 1H** *p* = 0.02) and phenylalanine metabolism (**Fig. 1I** *p* = 0.006) were significantly increased. This is of note as both pathways are implicated in gut-brain signaling. A full list of pathways altered by genotype can be found in **Supplemental Table 2**.

Given that the production of neuroactive metabolites is one of the most likely gut-brain signaling pathways, targeted metabolomics analysis was performed on levels of short-chain fatty acids (SCFA) and amino acids from cecal contents of Wt and *Shank3^KO^* mice^31,32^. Analysis of the three main SCFA, acetate, propionate, and butyrate demonstrated a significant decrease specifically in acetate in *Shank3^KO^* (**Fig. 1J** – *p* = 0.03). Analysis of select amino acid metabolites revealed genotype differences in multiple amino acids including the essential amino acids leucine and phenylalanine (**Fig. 1K**).

### Antibiotic throughout development exacerbates microbial, metabolic, and behavioral phenotypes in *Shank3^KO^* mice

To further interrogate the effect of the gut microbiome in *Shank3* deficient mice, a series of experiments using antibiotic-induced microbiome depletion (*<u>Abx</u>*) starting immediately after weaning (~P21) and continuing through early adulthood (~P60 – **Fig. 2A**) were conducted. By robustly reducing microbiome bulk and diversity, the full extent of interaction between *Shank3* genotype and microbiome status can be assessed. Importantly, this intervention did not affect body weight gain over time (**Fig. 2B**; Main effect of time: F_(1.44, 24.40)_ = 260.0 - *p*<0.0001) or levels of drinking across the experiment (**Fig. 2C**; *p* = 0.35). As expected, Abx treatment markedly reduced diversity of the microbiome using alpha (**Fig. 2D**; Main effect of Abx - F_(1,16)_ = 19.82; *p*=0.0004) and beta (**Fig. 2E**) diversity measures. From our targeted metabolomics analysis levels of acetate are affected by microbiome depletion (Main effect of Abx: F_(1,17)_ = 12.67; *p*=0.002) and that this effect is particularly pronounced in *Shank3^KO^* mice (**Fig. 2F** – Main effect of genotype: F_(1,17)_ = 9.615; *p*=0.007).

To test how manipulations to microbiome composition would affect *Shank3^KO^* behavioral phenotypes, control and antibiotic treated mice from both genotypes were assessed on a series of well-established ASD relevant behavioral paradigms. First, on a three chamber social interaction task there was no main effect of Group (F_(3,108)_ = 0.21; *p*=0.89) or Social Stimulus (F_(1,108)_ = 0.52; *p*=0.47), however there was a strong Group by Interactor interaction (**Fig. 2G -** F_(3,108)_ = 6.35; *p*=0.0005). Pairwise post-hoc testing demonstrates that Wt mice form the expected preference for the mouse interactor chamber vs the control cup (**2G light blue bars** - *p* = 0.0046), however Wt mice treated with Abx did not express significant social preference (**2G dark blue bars -** *p* = 0.4). *Shank3^KO^*-H_2_O mice showed no baseline social preference as expected (**2G pink bars** - *p* = 0.4), and *Shank3^KO^*-Abx mice showed a significant aversion for the socially paired chamber (**2G red bars** – *p* = 0.01). Thus, demonstrating that both microbiome manipulation and *Shank3* deletion impair sociability, with stronger effects in microbiome depleted *Shank3^KO^* mice.

Similarly treated mice were then assessed on an open field task. On a ten-minute test there was no effect on distance travelled (**Fig. 2H** – All main effects and interactions *p* > 0.35). When open field activity was assessed for 60 minutes there was a marked effect of genotype (**Fig. 2I** – Main effect of genotype: F_(1,57)_ = 11.15; *p*=0.002) consistent with previous reports on this mouse line^33^. On this measure there was no significant effect of time spent in the center (**Fig. 2J** – All p > 0.14) – which is frequently used as a measure of anxiety-like behavior. Anxiety-like behavior was further assessed using an elevated zero maze where there were no significant differences with genotype (**Fig. 2K** - F_(1,53)_ = 2.7; *p* =0.11) or treatment (F_(1,53)_ = 2.99; *p*=0.09). However, *Shank3^KO^* mice did show a modest increase in the number of total open arm entries (**Fig. 2L** - F_(1,54)_ = 6.61; *p* =0.013). Taken together, we find no consistent effect of *Shank3* deletion or microbiome depletion on anxiety like behavior, and genotype effects on locomotion when assessed over a longer period.

### Microbiome depletion interacts with *Shank3* genotype to affect transcriptional regulation in medial prefrontal cortex

Following behavioral testing, mice were rapidly decapitated and medial prefrontal cortex (mPFC) was dissected for full transcriptomic profiling via RNA sequencing (**Fig. 2A**). This brain region was selected as it is known to be critical for mediating social behaviors^34–36^, is directly affected by *Shank3* deletions^37–39^, and is affected by alterations in microbiome composition^7,40,41^.

Wt mice treated with Abx had significant differential expression of 522 genes relative to Wt-H_2_O controls (**Fig. 3A** and **Supp table 3**), and pathway analysis demonstrated that there were significant upregulations in genes related to mRNA translation and intracellular anatomical structures, and downregulation of pathways related to synaptic signaling and protein folding among others (**Fig. 3B**). KO mice on control water had differential expression of a similar number of genes (329 – **Fig. 3C**) with downregulation of pathways related to regulation of transcription and metabolic processes (**Fig. 3D**). Interestingly, the interaction between genotype and microbiome treatment was not as robust as the behavioral effects seen in Fig. 2G. There were fewer total regulated genes and less robust pathway mapping in KO-Abx mice than either of the previous conditions (**Fig. 3E/F**). All differentially regulated genes and pathway analyses are available as **Supplemental Tables 3-5**.

**Figure 3:**
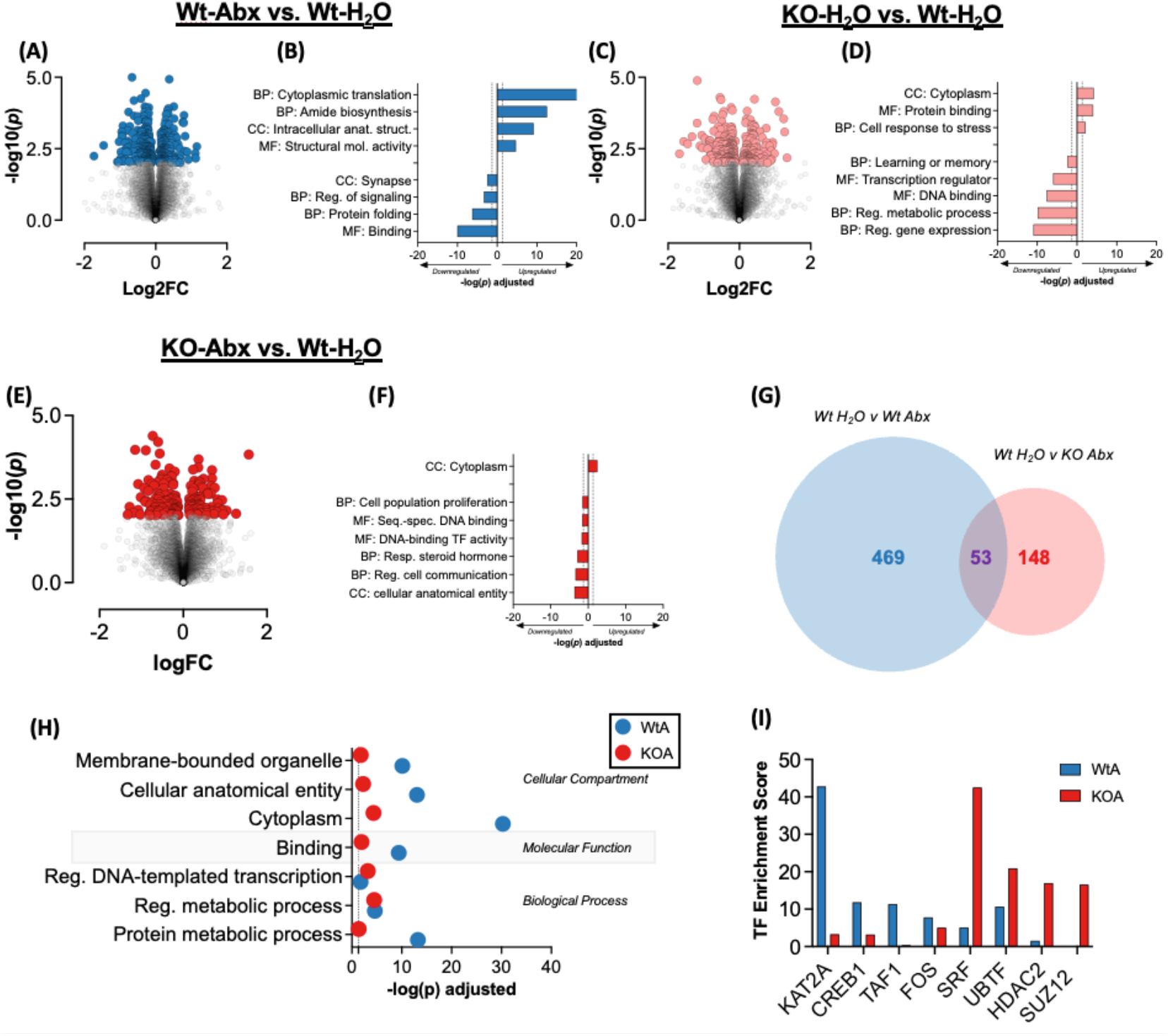
Effects of Shank3 Deletion and Antibiotic Treatment on mPFC Gene Expression. **(A)** Volcano plot of gene expression changes comparing Wt-H_2_O and Wt-Abx. **(B)** Select gene pathways regulated in Abx treated Wt mice. Dotted line at 1.3 indicates significance (FDR corrected *p*<0.05). **(C)** Volcano plot of gene expression changes comparing Wt-H_2_O and Shank3^KO^-H_2_O mice. **(D)** Select pathways that are predicted to be regulated in Shank3^KO^ mice. **(E)** Volcano plot of gene expression changes comparing KO-H_2_O and KO-Abx. **(F)** Select pathways that are predicted to be regulated in KO-Abx mice. **(G)** Venn diagram of all significant genes in Wt-Abx and KO-Abx relative to Wt-H_2_O controls shows the two genotypes have highly disparate responses to prolonged microbiome depletion. **(H)** Gene ontology analysis of regulated genes in Wt and KO mice following Abx. **(I)** Top Transcription Factors (TF) predicted to be upstream from differentially expressed genes in Wt-Abx and KO-Abx groups relative to Wt-H_2_O controls. *n* = 3-6 mice per group. Both male and female mice used in equal ratios.

**Figure 4:**
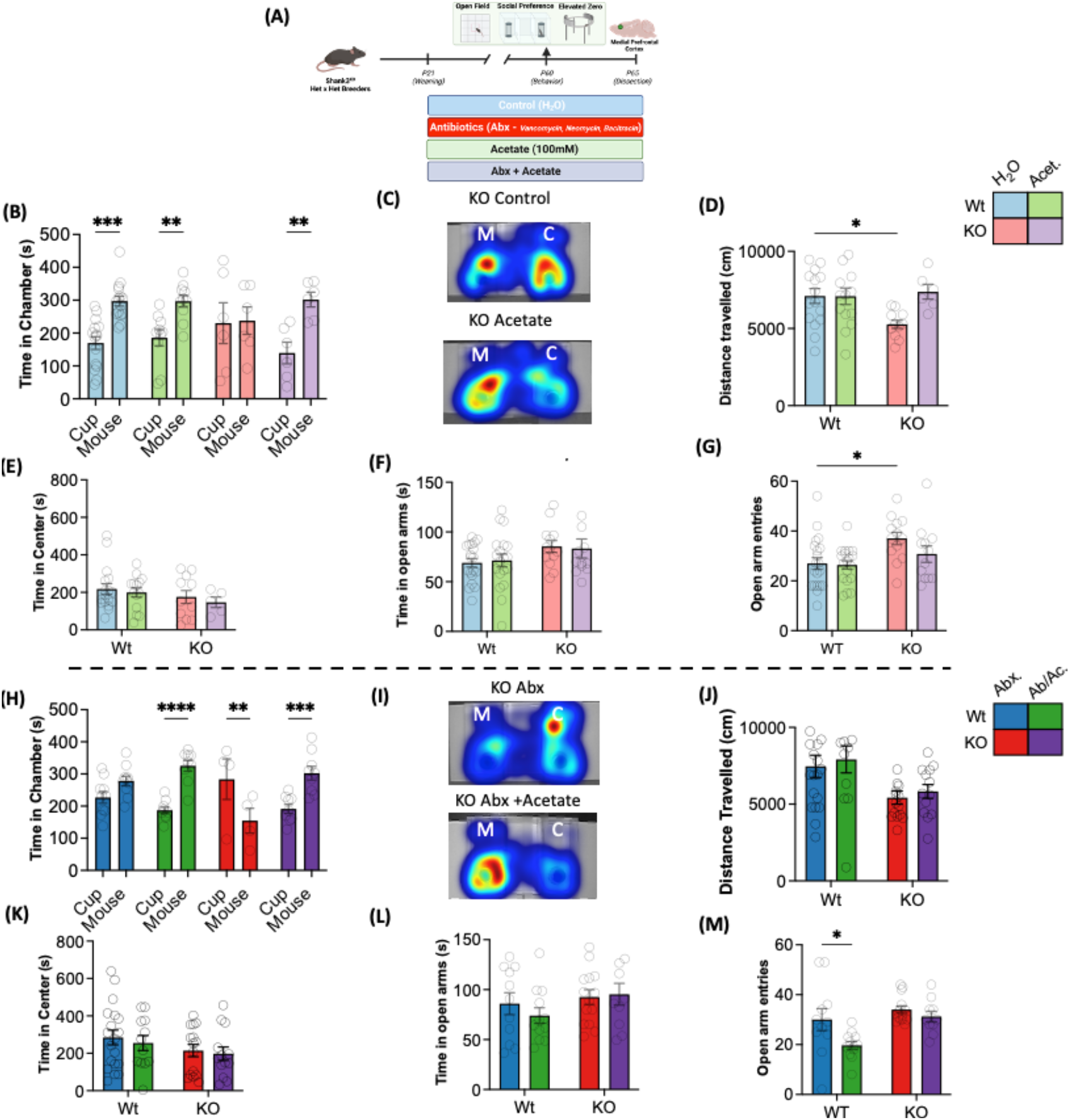
Acetate Replenishment Rescues Social Deficits Caused by Shank3 Deletion Independent of an Intact Microbiome. **(A)** Study design for Experiment 2 and mPFC collection. **(B)** In the three-chambered social interaction task, Wt-H_2_O and Wt-Acetate mice spent increased time in the chamber containing the novel social interactor (blue and green bars). Shank3^KO^-H_2_O did not show a significant preference for the social interactor at baseline as previous (pink bars). Acetate treatment rescues this social deficit in Shank3^KO^ mice (purple bars). **(C)** Representative Heatmap showing acetate treatment causes Shank3^KO^ mice to spend more time in Mouse (M) chamber (Bottom). **(D)** Acetate treatment also reverses locomotor activity deficit in Shank3^KO^ on one-hour open field task. **(E)** No genotype or treatment effects on time spent in center of open field in a 1-hour test session. **(F)** Shank3 ^KO^ have main effect of genotype in time spent in open arms of elevated zero maze with no interaction with acetate. **(G)** Shank3 ^KO^-H_2_O mice display increased frequency of entries into the open arm, but not those treated with acetate. **(H)** In the three-chambered social interaction task, Wt-Abx mice do not show a significant preference for the novel social interactor (blue bars). Treatment with acetate in microbiome depleted (Ab/Ac) mice reverses this phenotype (green bars). Shank3 ^KO^-Abx mice again demonstrate an aversion for the social stimulus (red bars), but this is reversed when acetate is supplemented to the antibiotics (Ab/Ac – purple bars). **(I)** Representative heatmap showing Ab/Ac treatment causes Shank3^KO^ to spend more time in the Mouse (M) chamber (Bottom). **(J)** Abx or Ab/Ac do not alter Shank3 genotype effects on locomotion in a 1-hour open field test session. **(K)** No genotype or treatment effects on time spent in center of open field in a 1-hour test session. **(L)** Treatment with Abx or Ab/Ac does not alter Shank3 genotype effects on time spent in open arms of the o-maze. **(M)** Wt Ab/Ac mice display decreased frequency of entries into the open arm of the elevated O-maze compared to Wt-Abx mice. All data presented as mean ± SEM. \**p* < 0.05; \*\**p* < 0.01; \*\*\**p* < 0.001; \*\*\*\**p* < 0.0001. *n* = 6-19 mice per group. Both male and female mice used in equal ratios.

When examined in more granular detail, there were marked contrasts between gene expression patterns induced by Abx treatment in Wt and KO mice. When the differentially expressed gene lists were compared, very little overlap was found, with distinct gene expression signatures after antibiotics in each genotype (**Fig. 3G**). Gene ontology analysis of the unique Abx-regulated genes in each genotype found that effects on pathways related to protein metabolism, binding, and cellular anatomy were markedly more affected in Wt-Abx mice than in *Shank3^KO^*-Abx mice (**Fig. 3H**). Similarly, when Enrichr was used to identify transcriptional regulators and epigenetic enzymes involved in the two gene sets, there are significantly different predicted regulators following Abx treatment in the two genotypes (**Fig. 3I**). Taken together, these findings suggest that *Shank3* genotype significantly affects the mPFC transcriptional response to microbiome depletion and may provide evidence for the discrepant behavioral phenotypes.

### Acetate Replenishment Rescues Social Deficits Caused by *Shank3* Gene Deletion Independent of Microbiome Status

Given findings of disrupted microbiome composition and function in *Shank3^KO^* mice (**Fig. 1**), decreased sociability that is exacerbated by microbiome depletion (**Fig. 2**), and significant effects of both *Shank3* knockout and microbiome depletion on gene expression in the mPFC (**Fig. 3**), was next investigated a potential role for gut-derived metabolites that may account for these phenotypes As shown above, levels of the SCFA acetate are decreased in the cecum of *Shank3^KO^* mice, and further diminished by Abx treatment. SCFA’s are a class of molecules produced almost exclusively by the gut microbiome, secondary to fermentation of insoluble fibers, and have been shown to play numerous roles in gut brain signaling^25^. Acetate in particular is known to inhibit histone deacetylase activity^42^, alter gene expression in the brain and behavioral responses to alcohol^43^, and play a role in regulating central satiety mechanisms^44^. Given such evidence, manipulations to levels of acetate were performed in *Shank3^KO^* mice followed by behavioral assessment to determine its mechanistic function.

For these experiments, both Wt and KO mice were put on the same treatments and time course as experiments described above, with additional groups treated with Acetate only or Abx+Acetate (Ab/Ac) (**Fig. 4A**). To assess effects of Acetate supplementation without microbiome manipulation on behavior, control water treated groups were directly compared to Acetate groups. Similarly, to determine the effect of acetate in reversing effects of microbiome depletion, Abx groups were directly compared to Ab/Ac groups. Importantly, Acetate and Ab/Ac treatment did not affect body weight gain overtime (**Supp Fig. 1A** Main effect of time: F_(1,29)_ = 253.0 - *p*<0.0001) or levels of drinking across the experiment (**Supp Fig. 1B** *p*>0.1). Treatment with Abx again markedly reduced diversity of the microbiome when compared to controls using beta diversity (**Supp Fig 1C)** and alpha diversity (**Supp Fig 1D and 1E**: Simpson Main effect of Abx F_(1,18)_ = 59.40 - *p*<0.0001 and Shannon Main effect of Abx F F_(1,18)_ = 658.7 - *p*<0.0001 respectively). Interestingly, treatment with acetate interacts with genotype to influence microbiome diversity (**Supp Fig 1F and 1G**: Simpson Interaction F_(1,17)_ = 7.734 - *p*<0.01) and Shannon Interaction F_(1,17)_ = 8.927 - *p*<0.01 respectively) when compared to control animals. As expected, treatment with acetate did not reverse Abx effects on microbiome diversity (**Supp Fig 1H and 1I** *p* >0.3).

In the three-chamber social interaction paradigm, treatment with Acetate resulted in a main effect of social stimulus (**Fig. 4B** - F_(1,66)_ = 25.90; *p* < 0.0001) no significant main effect of group (F_(3,66)_ = 0.17; *p* =0.91) and a borderline social stimulus by group interaction (F_(3,66)_ = 2.18; *p* =0.09). Planned pairwise Holm-Sidak post hoc comparisons showed Wt-H_2_O mice form a significant preference for the social stimulus (**3B** blue bars – *p* = 0.0002) which was not affected by acetate supplementation (**4B** green bars – *p* = 0.006). As with our previous experiments, KO-H_2_O mice did not form a significant preference for the social stimulus (**4B** pink bars – *p* = 0.87). However, KO-Acetate mice showed a marked reversal of the *Shank3* genotype effect and exhibit a significant preference for the social chamber (**4B** purple bars – *p* = 0.003). Similar effects of acetate were seen on distance traveled in a 1-hour open field task where there were no main effects of treatment or genotype (*p* > 0.05 for both), but there was a significant genotype by treatment interaction (F_(1,43)_ = 4.11; *p* =0.04). Post hoc testing shows that while the KO-H_2_O mice still exhibit the hypomobile phenotype, this effect is reversed by acetate supplementation (**Fig. 4D**).While there were no significant effects of genotype or treatment on time in the center of the open field (**Fig. 4E** - *p* > 0.17 for all), further assessment of anxiety-like behavior using the elevated zero maze, showed a modest genotype effect on time in the open arms (**Fig. 4F** - F_(1,59)_ = 4.69; *p* =0.03), but no significant effect of treatment or an interaction (*p* > 0.74 for both). As seen in our initial cohorts, the frequency of open arm entries was increased by *Shank3^KO^* status (**Fig. 4G** - F_(1,60)_ = 8.70; *p* =0.0045). Interestingly, Post hoc pairwise comparison showed only KO-H_2_O mice displayed increased open arm entries compared to Wt-H_2_O (*p* = 0.013), but not KO-Acetate mice– suggesting that acetate treatment may serve to moderate this behavioral phenotype as well.

Next, the effects of acetate supplementation on the behavioral effects of microbiome depletion in Wt and *Shank3^KO^* mice was assessed. In the three chambered social interaction task, treatment with Ab/Ac resulted in a main effect of interactor (F_(1,56)_ = 7.26; *p* =0.009), no main effect of group (F_(3,56)_ = 0.87; *p* =0.46), and a strong interactor by group interaction (**Fig. 4H** - F_(3,56)_ = 11.84; *p* <0.0001). As seen in the initial cohort, Wt mice treated with Abx did not exhibit significant preference for the social stimulus (**4**H blue bars – *p* = 0.1) and KO-Abx mice exhibited a significant aversion of the social chamber (**4H** red bars – *p* = 0.004). However, when mice were treated with antibiotics in combination with acetate, Abx induced social deficits were completely reversed in both Wt (**4H** green bars – *p* < 0.0001) and KO (**4H** purple bars – *p* = 0.0003) mice. Importantly, serum metabolomics show a main effect of acetate treatment in increasing serum acetate with minimal other effects on the metabolome (**Supplemental Figure 2**)

In the open field task, Ab/Ac treatment did not obviate the genotype effect of decreased locomotion at one hour (**Fig. 4J** - F_(1,55)_ = 9.51; *p* =0.003), and there were no main effects or interactions on time in center of the open field (**Fig. 4K**). When anxiety-like behavior was assessed on the elevated zero maze there were no main effects or interactions on time in open arms (**Fig. 4L**), but there was a significant effect of genotype in increasing number of open arm entries (**Fig. 4M** - F_(1,43)_ = 9.26; *p* =0.004) as well as a main effect of treatment on reducing the number of open arm entries (F_(1,43)_ = 6.62; *p* =0.01).

Taken together, these behavioral phenotypes suggest that acetate, which is reduced in *Shank3^KO^* and further reduced by antibiotics, robustly reverses social deficits caused by *Shank3* genotype, microbiome status, and the interaction. Additionally, acetate supplementation has some more modest effects on microbiome diversity and other behavioral measures including hypolocomotion and anxiety-like behaviors.

### Treatment with acetate induces unique transcriptional patterns in the mPFC

RNA-sequencing on the medial prefrontal cortex was next conducted on mice from the acetate supplementation study, to identify effects of acetate on central transcriptional regulation. Prolonged treatment with Acetate or Ab/Ac resulted in robust changes in gene expression in all groups relative to Wt controls (**Fig. 5A-D**). Given the ability of acetate treatment to rescue *Shank3^KO^* genotype effects on social behavior, we examined in detail the similarity in gene expression patterns between the two groups following treatment. Here, we see that there is considerable overlap in the significantly regulated genes in Wt and *Shank3^KO^* mice treated with acetate, but there are hundreds of uniquely regulated genes between the two genotypes (**Fig. 5E**). When the fold change of these individual genes was assessed, it is seen that the genes in common between the two lists were all changed in the same direction (**Fig. 5F** – Sig Both), and that most genes significantly regulated in one genotype had the same direction of change in the other genotype (**5F** – Sig Wt/KO). Similar patterns of gene overlap and directionality were seen in mice treated with the combination Ab/Ac treatment (**Fig. 5G-H**).

**Figure 5:**
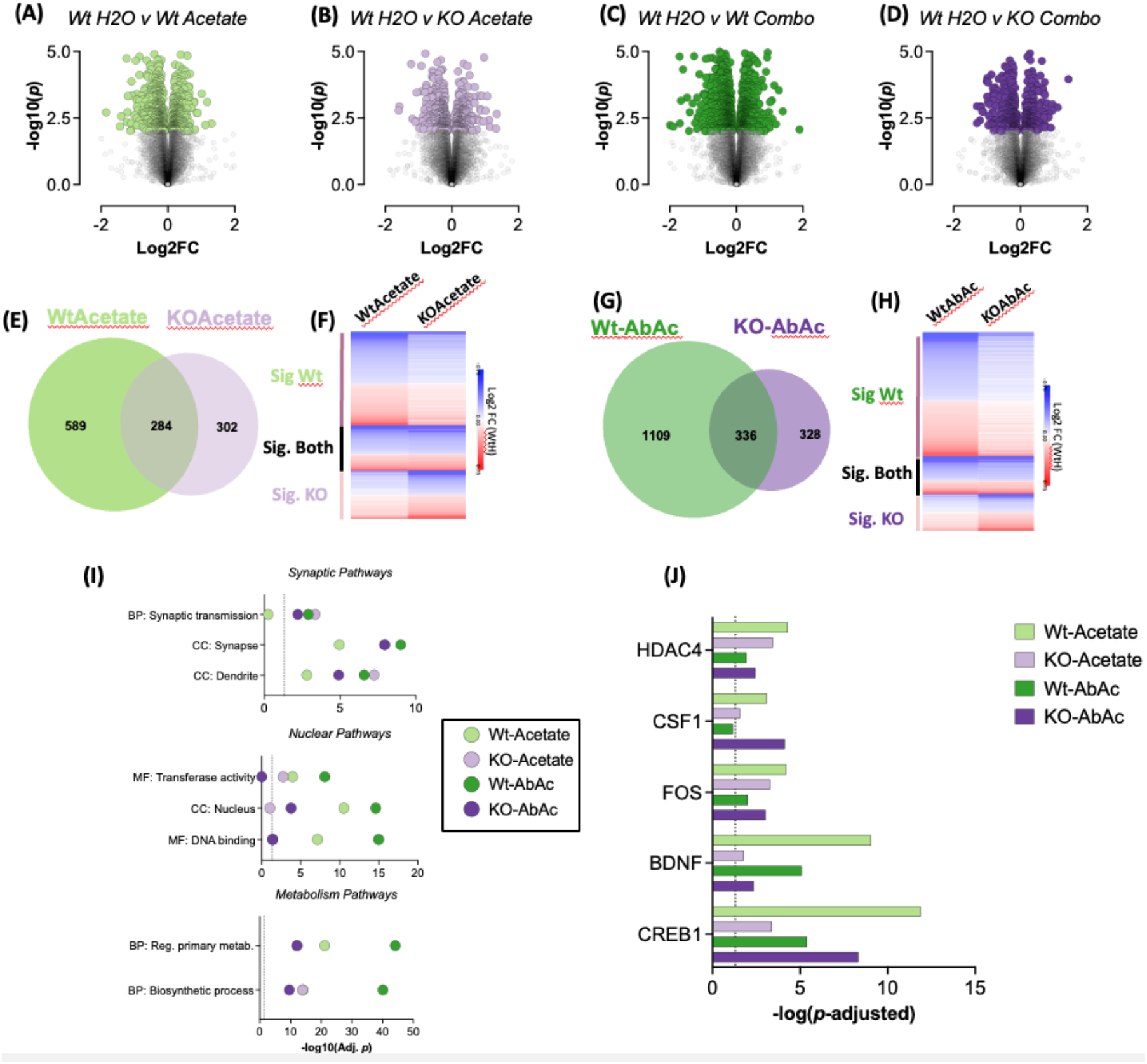
Acetate Replenishment With or Without Antibiotic Treatment on mPFC Gene Expression. **(A-D)** Volcano plots of all Acetate and Ab/Ac treatment groups relative to Wt-H_2_O controls. (**E)** Venn diagram of all genes significantly regulated in Wt-Acetate and KO-Acetate relative to Wt-H_2_O. **(F)** Heatmap of fold change expression of all genes from the previous panel. **(G)** Venn diagram of all genes significantly regulated in Wt-Ab/Ac and KO-Ab/Ac relative to Wt-H_2_O. **(H)** Heatmap of fold change expression of all genes from the previous panel. **(I)** Select pathways predicted to be enriched in uniquely regulated genes in Wt and KO mice from the four treatment groups. **(J)** Upstream regulator analysis of top enriched gene transcription regulators in Wt and KO mice from the four acetate treatment groups relative to Wt-H_2_O controls. *n* = 4-6 mice per group. Both male and female mice used in equal ratios.

To see how treatment with acetate or combination Ab/Ac affected gene regulation pathways G:Profiler pathway analysis was performed^1^. Treatment with acetate and Ab/Ac resulted in robust regulation of metabolic pathways in all groups with a more robust effect seen in Wt mice (**Fig. 5I** – bottom). A similar pattern is seen with nuclear and transcription related genes which were robustly significant in Wt mice, but only marginally so in *Shank3^KO^* (**Fig. 5I** – middle). Genes related to synaptic function were also differentially expressed in all groups but were generally more robustly changed in *Shank3^KO^* mice (**Fig. 5I** – top). Ingenuity pathway analysis of predicted upstream regulators of significantly regulated genes in each group found that numerous activity dependent transcription factors (Fos, Creb) and neurotrophic and gliotrophic factors (BDNF, Csf1) were predicted to be upstream of regulated genes in these four Acetate and Ab/Ac treated groups (**Fig. 5J**). Full pathway analyses from these comparisons are available in **Supplemental Tables S5 & S8**.

Given the complexity of our sequencing analysis between all groups, we also performed weighted gene coexpression network analysis (WGCNA)^45^. This analysis allows for data driven means of identifying networks of genes that cluster together based on expression in individual treatment types. **Fig. 6A** presents the dendrogram separating genes into modules. Associations of specific gene modules with genotype, sex, Abx or Acetate treatment were performed, and significant associations are highlighted in **Fig. 6B**. To obtain more detailed analysis of the gene expression effects of our treatments, we highlight multiple significant modules. The purple module was found to be negatively associated with both Abx and Acetate treatment. This association is highlighted on the z-scored heatmap of genes in the module, showing decreased expression of all purple module genes in all treatment groups (**Fig. 6C**). This module was significantly associated with nuclear processes including nucleic acid binding pathways, and RNA metabolic processing (**Fig. 6D**). STRING analysis was used to identify the functional connections between resultant gene products as shown in **Fig. 6E**, this produces a highly interactive network with interconnections far exceeding those predicted by chance. Strongly regulated pathways are highlighted in **Fig. 6F** cloud diagram.

**Figure 6:**
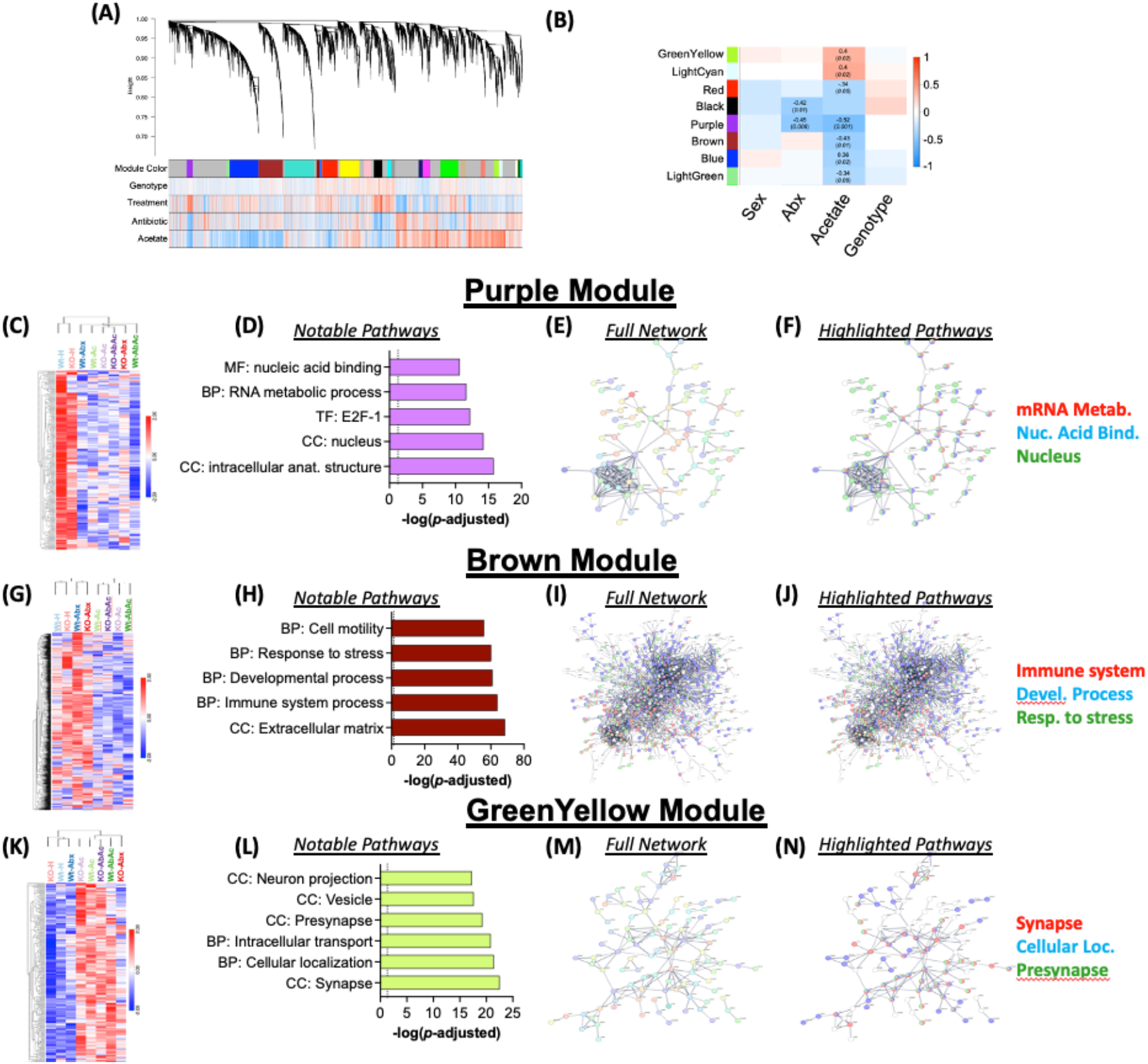
Weighted Gene Network Analysis Reveals Molecular Mechanisms Which correlate with Acetate Replenishment With or Without Antibiotic Treatment. **(A)** Dendrogram of WGCNA analysis. **(B)** Module-trait relationship table showing modules of interest. Each cell reports the Pearson correlation value and if significant the *p*-value in parentheses. Columns describe the variable and the rows show the module name **(C)** Heatmap showing unsupervised hierarchical clustering of all genes in Purple module. **(D)** Gene ontology enrichment of the differentially expressed genes in Purple module. **(E)** Protein-protein interaction (PPI) network enrichment for the Purple module from the STRING database. **(F)** Select pathways from full PPI network enrichment for the purple module from STRING database. **(G-J)** Heatmap, gene ontology enrichment and PPI network enrichment for genes in Brown module. **(K-N)** Heatmap, gene ontology enrichment and PPI network enrichment for genes in Green Yellow module. *n* = 4-6 mice per group. Both male and female mice used in equal ratios.

As acetate treatment was the most robustly associated with gene networks, we also provide detailed analysis of the Brown module which is negatively associated with acetate treatment and has significant pathways associated with immune system processes, developmental processes, and response to stress among others (**Fig. 6G-J**). The GreenYellow module which is positively associated with acetate treatment led to regulation of genes associated with synaptic pathways and cellular localization (**Fig. 6K-N**). Full pathway analyses from these modules is found in **Supplemental Table 9**. Taken together these data demonstrate robust transcriptional effects of acetate in key behaviorally relevant brain regions.

### Levels of Acetate in the serum of patients with Phelan McDermid Syndrome Correlate with ABC Hyperactivity Scores in a Sex Dependent Manner

To assess the relevance of the baseline changes in SCFA levels observed in the *Shank3^KO^* model to humans, we generated metabolomic data targeting levels of eight SCFA’s including acetate across a subset of 32 PMS probands (who are heterozygous for the *Shank3* gene) and their unaffected controls (**Fig.7A**). The main sociodemographic and clinical characteristics of study participants are shown in **Table S11**.

**Figure 7:**
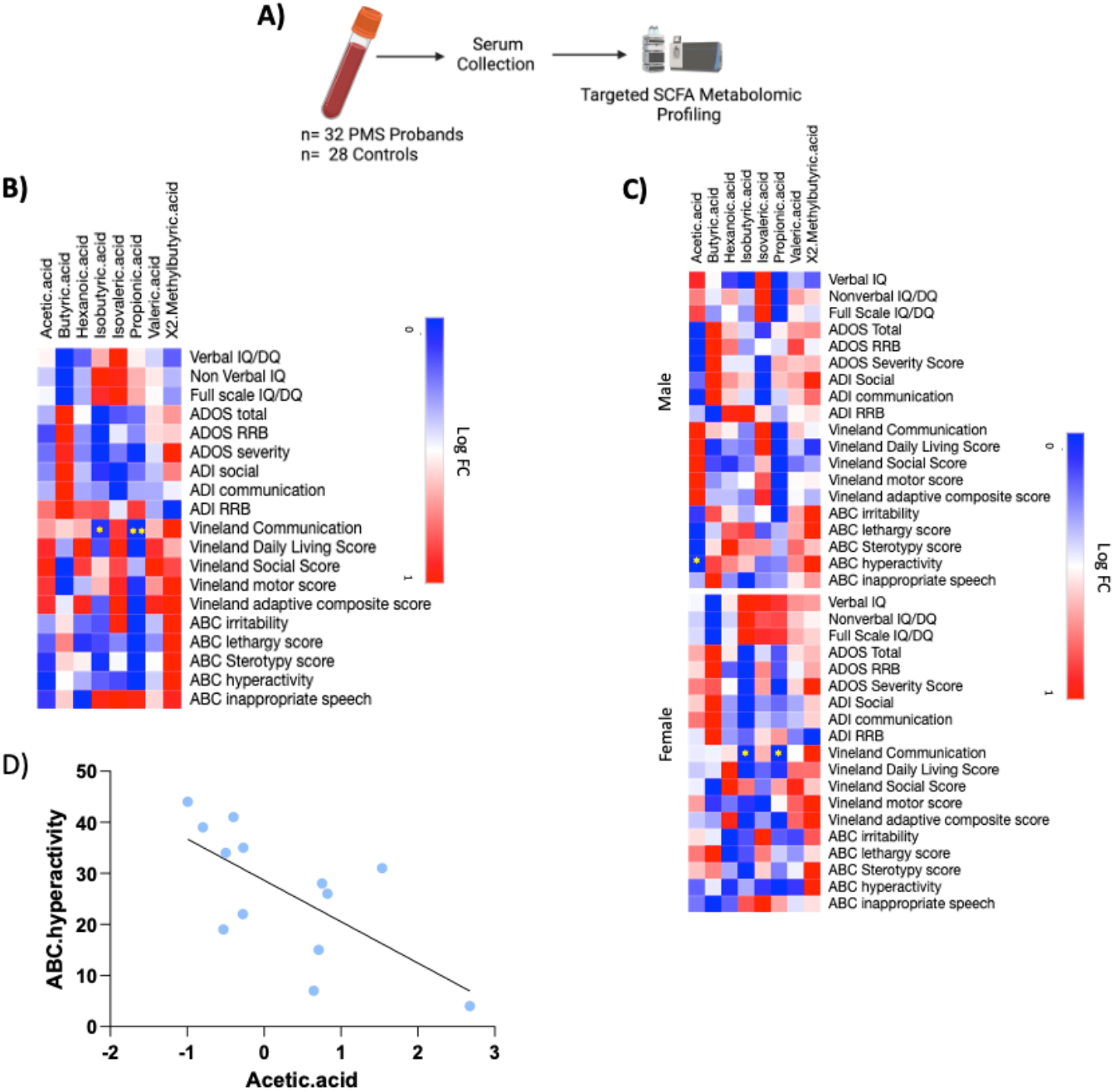
Serum Levels of Acetate Correlate with ABC Hyperactivity in Male PMS Patients. **(A)** Study Design: Clinical scores as well as serum was collected from 32 PMS probands and 28 unaffected family controls. Serum was then analyzed for levels of 8 SCFAs using liquid chromatography-mass spectrometry (LC/MS) (**B**) Heatmap showing a significant correlation between Vineland Communication scores and levels of Isobutyric acid and propanoic acid (n =30) (**C**) Heatmap showing a significant negative correlation between Vineland Communication scores and levels of Isobutyric acid and propanoic acid in females only (n =17) and significant negative correlation between ABC hyperactivity scores and Acetic Acid levels in males (n= 13) **(D)** Spearman’s correlation confirming significant negative correlation between levels of acetic acid and ABC hyperactivity in males (R= −0.5824). \**p* < 0.05; \*\**p* < 0.01

PCA analysis comparing PMS probands to controls showed PC1 explained 40.87% of variance in the data (**Fig.S4A**). Next, we conducted a primary analysis to look for the effect of genotype (PMS or control) variant class (mutation or deletion) variant type (frameshift, ring complex or simple deletion) as well as size of deletion on circulating levels of SCFA (**Fig. S4B**). We report no significant effect of any of the above variables on serum levels of SCFA’s, not even when the analysis is conducted in a sex dependent manner (**Fig. S4C**). Subsequently, a secondary analysis was carried out where we modelled for the effect of several clinical variables on the differential abundance of SCFAs as described below. Interestingly our results show levels of Isobutyric acid and Propionic acid negatively correlated with Vineland communication scores (**Fig. 7B** *n*= 30 Adjusted *p*<0.05 and *p*<0.01 respectively). Sex specific analysis further showed the negative correlation between Vineland communication scores and both levels of Isobutyric acid, Propionic and was driven by female PMS patients (**Fig. 7C** *n*= 17 Adjusted *p*<0.05). Moreover, levels of acetate were found to show a significant negative correlation with ABC hyperactivity scores in males only (**Fig. 7C** n = 13 Adjusted *p*<0.05). A Spearman’s correlation confirmed the significant negative correlation between levels of acetate and ABC hyperactivity in males (R= −0.58 p<0.05; **Fig. 7D).** Taken together, clinical findings strengthen the hypothesis that targeting altered levels of SCFAs in PMS patients may help improve specific clinical measures including communication and hyperactivity.

## Discussion

Here we demonstrate that mice lacking all isoforms of the *Shank3* gene have marked disruptions of gut microbiome composition and function at baseline (**Fig. 1**). Further depletion of the microbiome with antibiotics leads to reduction in social preference in Wt mice, and further exacerbates the decreased social preference of *Shank3^KO^* mice (**Fig. 2**). Microbiome depletion is also associated with dysregulation of the transcriptome of the frontal cortex in both Wt and *Shank3^KO^* mice (**Fig. 3**). Importantly, we find that *Shank3^KO^* mice are deficient in the microbial metabolite acetate, and this deficiency is further exacerbated by antibiotic treatment (**Fig. 2F**). Oral treatment with acetate reverses both the *Shank3* genotype effects, and microbiome depletion effects in Wt and *Shank3^KO^* mice on the social interaction task (**Fig. 4**), and reverses genotype effects on hypolocomotion with modest effects on anxiety like behaviors in *Shank3^KO^*. Mice treated with acetate show marked transcriptional changes in the medial prefrontal cortex of both genotypes, with marked effects on pathways related to synaptic signaling, DNA binding, and metabolism (**Fig. 5**). This was particularly evident given robust acetate responsive modules of gene coexpression identified by WGCNA (**Fig. 6**). Finally, utilizing samples from a clinical population, we verify that there are correlations between levels of acetate and specific behavioral phenotypes in patients with Phelan-McDermid Syndrome – a condition driven by mutations in the *Shank3* gene (**Fig. 7**). Together these findings provide additional important mechanistic evidence for gut-brain signaling in ASD.

### Dysregulation of microbiome composition in *Shank3^KO^* mice

We find that *Shank3^KO^* mice display a significant decrease in bacterial diversity of their gut microbiome with a decrease in the ratio of the two major phyla of bacteria, Firmicutes and Bacteroidetes (**Fig. 1**). Greater than 90% of the mammalian microbiome is comprised of bacteria from the Firmicutes and Bacteroidetes phyla, and changes in the ratio of these phyla are thought to be an important marker of microbiome dysregulation^47^. Shifts in this ratio are associated with behavioral response to social stress^48^ as well as with changes in production of the important short chain fatty acid metabolites^49^. A recent study utilizing a *Shank3* mutant mouse lacking both the α and β isoforms of *Shank3* also reported a shift in the ratio of Firmicutes to Bacteroidetes^50^. However, while this group did find a similar decrease in bacterial diversity, they found the opposite change in Firmicutes/Bacteroidetes ratio. Similarly, human studies report some patients with ASD present with a decreased Firmicutes/Bacteroidetes ratio^51,52^ while others show the opposite result^53,54^.

At the genus level we report a decrease in Lactobacillus bacteria across all phylogenetic levels measured (**Fig. 2G**). This is in line with two additional studies that utilized a model of *Shank3* B deletion and report specific decreases in bacteria from the *Lactobacillus* genus in *Shank3* B knockout mice^26,27^. Additionally, these studies noted that supplementation with the *Lactobacillus* species *L. reuteri* was able to rescue some level of social deficits in the *Shank3* B knockout mice. Changes in Lactobacillus were not noted in the report on *Shank3* A/B mice^50^. Together, our studies lend strength to evidence supporting reductions in *Lactobacillus* due to *Shank3* mutations and begin to suggest this bacterial group may be an important component of gut-brain signaling in *Shank3* models of ASD.

### Changes in bacterial metabolites and functional pathways in *Shank3^KO^*

The gut microbiome is a highly metabolically active ecosystem that produces hundreds of metabolites that are absorbed into circulation^56^. Some of the strongest evidence for gut-brain signaling mechanisms is via bacterial production of neuroactive metabolites^57^. The most widely studied class of these metabolites are the SCFA’s, which are produced by bacterial fermentation of fiber. The three most abundant of these – acetate, butyrate, and propionate - have been shown to be critical for regulation of blood-brain barrier integrity, microglial function, behavioral response to drugs of abuse, and epigenetic regulation in the brain^21,22,43,58^. We found that *Shank3^KO^* mice have marked decreases in the SCFA acetate, an effect that was exacerbated by antibiotic treatment (**Figs. 1J, 2F**). Acetate is known to function as a histone deacetylase inhibitor and it serves as a substrate for acetyl-CoA which is necessary for histone acetylation, thus giving acetate the potential to serve as an important epigenetic regulator. Indeed, recently published work has demonstrated that gut-derived acetate can affect levels of histone acetylation in the brain in behaviorally significant ways^43^.

In addition to changes in SCFA, *Shank3^KO^* mice also exhibited increases in levels of multiple amino acids (**Fig. 1K**). While functional consequences of these amino acid changes in the pathophysiology of ASD is not fully clear, there are amino acids of note from our findings. *Shank3^KO^* leads to an increase in levels of phenylalanine, increases in phenylalanine due to inborn errors of metabolism are well known to produce autism-like behavioral symptoms amongst a constellation of other symptoms in patients with phenylketonuria^59^. While less well studied, changes in serum levels of phenylalanine have also been reported in patients with idiopathic ASD^60^.

Moreover, KEGG Orthology metagenomic analysis confirmed an increase in Fatty Acid degradation (**Figs. 1H)** and phenylalanine metabolism (**Figs. 1I)** which likely corresponds to the decrease in levels of acetate and increase in Phenylalanine measured at the metabolite level. Together these findings demonstrate a robust change in composition and functional status of the microbiome due to *Shank3* gene deletion.

### ASD-like behavioral phenotypes and metabolic profiles are exacerbated by microbiome depletion and rescued by treatment with acetate in *Shank3^KO^*

For these studies we measured effects of genotype and microbiome targeted interventions on social behavior using the three chambered social interaction task. We find that *Shank3^KO^* mice have baseline deficits in social behavior, and that depletion of the microbiome with antibiotics results in mice of both genotypes displaying decreased social preference (**Fig. 2G**). In fact, the *Shank3^KO^*-Abx mice exhibited a significant aversion for the socially paired chamber. This increased behavioral sensitivity to microbiome depletion in KO mice is further corroborated by metabolomic analysis of serum showing KO mice had a more exaggerated response to Abx treatment with 51 significantly altered metabolites compared to 19 metabolites in Wt Abx treated mice (**Supp Fig. 2C**). KO-Abx mice also showed enrichment of the nicotinate and nicotinamide metabolism pathway (**Supp Fig. 2D**) a pathway highly implicated in a range of neuropsychiatric disorders. Given that both *Shank3* knockout and antibiotic treatment reduced the SCFA acetate, we set out to determine if supplementation with acetate via the drinking water could ameliorate behavioral effects. Indeed, we find that acetate treatment reverses the decreased sociability seen in *Shank3^KO^* mice with an intact microbiome (**Fig. 4B,C**). Treatment with acetate also reversed the decreases in sociability due to microbiome depletion in both Wt and *Shank3^KO^* mice (**Fig. 4H,I**) as well as normalizing Abx effects on serum metabolic profile, albeit to a larger extent in KO mice compared to Wt counterparts (**Supp Fig. 2C**). These findings are particularly notable given that treatment with a single metabolite takes *Shank3^KO^* Abx treated mice who have a baseline aversion for the social stimulus and returns it to a robust preference. These effects of microbiome manipulations on sociability are not surprising and there is a growing literature showing microbiome effects on social behaviors^17,19,35^, however these are among the first findings to demonstrate that these broad microbiome effects can be reversed via treatment with a single microbiome derived metabolite.

### Differential effects of microbiome depletion and acetate supplementation on cortical gene expression in Wt and *Shank3^KO^* mice

Previous reports have demonstrated that mice born germ free or with their microbiome altered by antibiotics have differential gene expression in the brain^7,9,21^. To assess how *Shank3* gene deletion might alter this, we performed RNA-sequencing of the medial prefrontal cortex (PFC). This region was chosen as it is known regulator of social behavior^34^, and multiple previous studies have shown it to be sensitive to depletion of the gut microbiome^7,40,41^. In our antibiotic only treatment groups, we found that microbiome depletion led to distinct patterns of gene expression in the two genotypes, with more robust effects on metabolism, and cytoplasm related genes being seen in the Wt-Abx mice (**Fig. 3**). When mice were treated with acetate with or without antibiotics we found more robust changes in gene expression patterns in the brain (**Fig. 5A-D**) with more convergence in the number of genes that were regulated in the two genotypes and greater symmetry in the direction of gene expression changes (**Fig. 5E-H**). Acetate drove expression in pathways related to synaptic function, transcription and metabolism and many of the genes that were altered by acetate treatment are predicted to be downstream of key activity dependent transcription factors (**Fig. 5I-J**). Gene coexpression network analysis also provided granular insight into acetate sensitive gene networks and that acetate treatment drove changes related to immune system processes, cell motility, and synaptic function (**Fig. 6**). Previous work, largely from the Berger lab, has shown that acetate metabolism can lead to transcriptional regulation and behavioral change in multiple paradigms^43,61,62^. Additionally, acetate can act as a histone deacetylase inhibitor and indirectly influence transcriptional patterns through these mechanisms^42,44^.

### Clinical correlates of acetate in patients with Phelan-McDermid Syndrome

To examine the translational relevance of our findings from the *Shank3^KO^* mouse model, we also assessed serum levels of acetate in patients diagnosed Phelan-McDermid Syndrome, a condition caused by mutations in the *Shank3* gene, compared to neurotypical controls. Despite not observing any genotype effects on levels of acetate, which may be due to the low n number of patients in our study. We observe that levels of acetate exhibit negative correlations with the ABC hyperactivity scale in male patients (**Fig. 6D**). These findings suggest a potential for use of acetate as a biomarker or therapeutic intervention in patients with ASD.

## Conclusions

Here we utilized a genetically valid model of ASD lacking both copies of the *Shank3* gene. We find that these mice have marked differences in the composition and functional output of their gut microbiome. Additionally, these mice were more susceptible to the behavioral metabolic and transcriptional effects of gut microbiome depletion. Importantly, these effects can be reversed by treatment with the gut-derived metabolite acetate. These experiments provide important additional data for a gene by microbiome interaction in the pathophysiology of ASD and have the potential to lead to bench to bedside translational research in ASD.

## Acknowledgements

This work was supported primarily by funds from the Seaver Autism Center and Friedman Brain Institute. K.R.M. is supported by NIH grants NS124187 & NS117356. K.L.C. was supported by a postdoctoral fellowship from the Canadian Institutes of Health Research (no. 201811MFE-414896-231226). L.L. is supported by National Institutes of Health (NIH) Medical Scientist Training Program T32 GM07170 and Training Grant in Computational Biology 5-T32-HG-000046–21.

## Author Contributions

A.O., N.L.M., A.N.S, T.J.E, G.Z., K.R.M., L.L., C.A.T. & D.D.K. performed experiments; A.O., S.R.T., K.L.C, A.B.G., E.D., L.L., J.G., C.A.T., M.S.B., & D.D.K analyzed data; A.O., E.D., J.D.B., M.S.B., & D.D.K contributed to experimental design; J.D.B. generated the mouse line; A.O. & D.D.K. wrote the manuscript; All authors contributed critical edits and feedback on the final manuscript draft.

## References

1. Volkmar, F.R., Reichow, B., and McPartland, J. (2012). Classification of autism and related conditions: progress, challenges, and opportunities. Dialogues Clin Neurosci 14, 229–237.

2. Maenner, M.J., Shaw, K.A., Bakian, A.V., Bilder, D.A., Durkin, M.S., Esler, A., Furnier, S.M., Hallas, L., Hall-Lande, J., Hudson, A., et al. (2021). Prevalence and Characteristics of Autism Spectrum Disorder Among Children Aged 8 Years - Autism and Developmental Disabilities Monitoring Network, 11 Sites, United States, 2018. MMWR Surveill Summ 70, 1–16. 10.15585/mmwr.ss7011a1.

3. Satterstrom, F.K., Kosmicki, J.A., Wang, J., Breen, M.S., Rubeis, S.D., An, J.-Y., Peng, M., Collins, R., Grove, J., Klei, L., et al. (2020). Large-Scale Exome Sequencing Study Implicates Both Developmental and Functional Changes in the Neurobiology of Autism. Cell 180, 568–584.e23. 10.1016/j.cell.2019.12.036.

4. Mila, M., Alvarez-Mora, M.I., Madrigal, I., and Rodriguez-Revenga, L. (2018). Fragile X syndrome: An overview and update of the FMR1 gene. Clin. Genet. 93, 197–205. 10.1111/cge.13075.

5. De Rubeis, S., Siper, P.M., Durkin, A., Weissman, J., Muratet, F., Halpern, D., Trelles, M.D.P., Frank, Y., Lozano, R., Wang, A.T., et al. (2018). Delineation of the genetic and clinical spectrum of Phelan-McDermid syndrome caused by SHANK3 point mutations. Mol Autism 9, 31. 10.1186/s13229-018-0205-9.

6. Cryan, J.F., and Dinan, T.G. (2012). Mind-altering microorganisms: the impact of the gut microbiota on brain and behaviour. Nat Rev Neurosci 13, 701–712. 10.1038/nrn3346.

7. Chu, C., Murdock, M.H., Jing, D., Won, T.H., Chung, H., Kressel, A.M., Tsaava, T., Addorisio, M.E., Putzel, G.G., Zhou, L., et al. (2019). The microbiota regulate neuronal function and fear extinction learning. Nature 574, 543–548. 10.1038/s41586-019-1644-y.

8. Luczynski, P., Whelan, S.O., O’Sullivan, C., Clarke, G., Shanahan, F., Dinan, T.G., and Cryan, J.F. (2016). Adult microbiota-deficient mice have distinct dendritic morphological changes: differential effects in the amygdala and hippocampus. Eur. J. Neurosci. 44, 2654–2666. 10.1111/ejn.13291.

9. Hoban, A.E., Stilling, R.M., Moloney, G., Shanahan, F., Dinan, T.G., Clarke, G., and Cryan, J.F. (2018). The microbiome regulates amygdala-dependent fear recall. Mol. Psychiatry 23, 1134–1144. 10.1038/mp.2017.100.

10. Adams, J.B., Johansen, L.J., Powell, L.D., Quig, D., and Rubin, R.A. (2011). Gastrointestinal flora and gastrointestinal status in children with autism--comparisons to typical children and correlation with autism severity. BMC Gastroenterol 11, 22. 10.1186/1471-230X-11-22.

11. Ashwood, P., Krakowiak, P., Hertz-Picciotto, I., Hansen, R., Pessah, I., and Van de Water, J. (2011). Elevated plasma cytokines in autism spectrum disorders provide evidence of immune dysfunction and are associated with impaired behavioral outcome. Brain Behav. Immun. 25, 40–45. 10.1016/j.bbi.2010.08.003.

12. McElhanon, B.O., McCracken, C., Karpen, S., and Sharp, W.G. (2014). Gastrointestinal symptoms in autism spectrum disorder: a meta-analysis. Pediatrics 133, 872–883. 10.1542/peds.2013-3995.

13. Macfabe, D.F. (2012). Short-chain fatty acid fermentation products of the gut microbiome: implications in autism spectrum disorders. Microb Ecol Health Dis 23. 10.3402/mehd.v23i0.19260.

14. Ahmed, H., Leyrolle, Q., Koistinen, V., Kärkkäinen, O., Layé, S., Delzenne, N., and Hanhineva, K. (2022). Microbiota-derived metabolites as drivers of gut-brain communication. Gut Microbes 14, 2102878. 10.1080/19490976.2022.2102878.

15. Stilling, R.M., Moloney, G.M., Ryan, F.J., Hoban, A.E., Bastiaanssen, T.F., Shanahan, F., Clarke, G., Claesson, M.J., Dinan, T.G., and Cryan, J.F. (2018). Social interaction-induced activation of RNA splicing in the amygdala of microbiome-deficient mice. Elife 7. 10.7554/eLife.33070.

16. Stilling, R.M., Ryan, F.J., Hoban, A.E., Shanahan, F., Clarke, G., Claesson, M.J., Dinan, T.G., and Cryan, J.F. (2015). Microbes & neurodevelopment--Absence of microbiota during early life increases activity-related transcriptional pathways in the amygdala. Brain Behav. Immun. 50, 209–220. 10.1016/j.bbi.2015.07.009.

17. Desbonnet, L., Clarke, G., Shanahan, F., Dinan, T.G., and Cryan, J.F. (2014). Microbiota is essential for social development in the mouse. Mol. Psychiatry 19, 146–148. 10.1038/mp.2013.65.

18. Fröhlich, E.E., Farzi, A., Mayerhofer, R., Reichmann, F., Jačan, A., Wagner, B., Zinser, E., Bordag, N., Magnes, C., Fröhlich, E., et al. (2016). Cognitive impairment by antibiotic-induced gut dysbiosis: Analysis of gut microbiota-brain communication. Brain Behav. Immun. 56, 140–155. 10.1016/j.bbi.2016.02.020.

19. Desbonnet, L., Clarke, G., Traplin, A., O’Sullivan, O., Crispie, F., Moloney, R.D., Cotter, P.D., Dinan, T.G., and Cryan, J.F. (2015). Gut microbiota depletion from early adolescence in mice: Implications for brain and behaviour. Brain Behav. Immun. 48, 165–173. 10.1016/j.bbi.2015.04.004.

20. Thion, M.S., Low, D., Silvin, A., Chen, J., Grisel, P., Schulte-Schrepping, J., Blecher, R., Ulas, T., Squarzoni, P., Hoeffel, G., et al. (2018). Microbiome Influences Prenatal and Adult Microglia in a Sex-Specific Manner. Cell 172, 500–516.e16. 10.1016/j.cell.2017.11.042.

21. Erny, D., Hrabě de Angelis, A.L., Jaitin, D., Wieghofer, P., Staszewski, O., David, E., Keren-Shaul, H., Mahlakoiv, T., Jakobshagen, K., Buch, T., et al. (2015). Host microbiota constantly control maturation and function of microglia in the CNS. Nat. Neurosci. 18, 965–977. 10.1038/nn.4030.

22. Kiraly, D.D., Walker, D.M., Calipari, E.S., Labonte, B., Issler, O., Pena, C.J., Ribeiro, E.A., Russo, S.J., and Nestler, E.J. (2016). Alterations of the Host Microbiome Affect Behavioral Responses to Cocaine. Sci Rep 6, 35455. 10.1038/srep35455.

23. Dohnalová, L., Lundgren, P., Carty, J.R.E., Goldstein, N., Wenski, S.L., Nanudorn, P., Thiengmag, S., Huang, K.-P., Litichevskiy, L., Descamps, H.C., et al. (2022). A microbiome-dependent gut–brain pathway regulates motivation for exercise. Nature 612, 739–747. 10.1038/s41586-022-05525-z.

24. Bachmann, C., Colombo, J.P., and Berüter, J. (1979). Short chain fatty acids in plasma and brain: quantitative determination by gas chromatography. Clin Chim Acta 92, 153–159. 10.1016/0009-8981(79)90109-8.

25. Dalile, B., Van Oudenhove, L., Vervliet, B., and Verbeke, K. (2019). The role of short-chain fatty acids in microbiota-gut-brain communication. Nat Rev Gastroenterol Hepatol 16, 461–478. 10.1038/s41575-019-0157-3.

26. Tabouy, L., Getselter, D., Ziv, O., Karpuj, M., Tabouy, T., Lukic, I., Maayouf, R., Werbner, N., Ben-Amram, H., Nuriel-Ohayon, M., et al. (2018). Dysbiosis of microbiome and probiotic treatment in a genetic model of autism spectrum disorders. Brain, Behavior, and Immunity 73, 310–319. 10.1016/j.bbi.2018.05.015.

27. Sgritta, M., Dooling, S.W., Buffington, S.A., Momin, E.N., Francis, M.B., Britton, R.A., and Costa-Mattioli, M. (2019). Mechanisms Underlying Microbial-Mediated Changes in Social Behavior in Mouse Models of Autism Spectrum Disorder. Neuron 101, 246–259.e6. 10.1016/j.neuron.2018.11.018.

28. Drapeau, E., Riad, M., Kajiwara, Y., and Buxbaum, J.D. (2018). Behavioral Phenotyping of an Improved Mouse Model of Phelan-McDermid Syndrome with a Complete Deletion of the Shank3 Gene. eNeuro 5. 10.1523/ENEURO.0046-18.2018.

29. Hills, R.D., Pontefract, B.A., Mishcon, H.R., Black, C.A., Sutton, S.C., and Theberge, C.R. (2019). Gut Microbiome: Profound Implications for Diet and Disease. Nutrients 11. 10.3390/nu11071613.

30. Fields, C.T., Sampson, T.R., Bruce-Keller, A.J., Kiraly, D.D., Hsiao, E.Y., and de Vries, G.J. (2018). Defining Dysbiosis in Disorders of Movement and Motivation. J. Neurosci. 38, 9414–9422. 10.1523/JNEUROSCI.1672-18.2018.

31. van Sadelhoff, J.H.J., Perez Pardo, P., Wu, J., Garssen, J., van Bergenhenegouwen, J., Hogenkamp, A., Hartog, A., and Kraneveld, A.D. (2019). The Gut-Immune-Brain Axis in Autism Spectrum Disorders; A Focus on Amino Acids. Front Endocrinol (Lausanne) 10, 247. 10.3389/fendo.2019.00247.

32. Dinan, T.G., and Cryan, J.F. (2017). The Microbiome-Gut-Brain Axis in Health and Disease. Gastroenterol. Clin. North Am. 46, 77–89. 10.1016/j.gtc.2016.09.007.

33. Drapeau, E., Riad, M., Kajiwara, Y., and Buxbaum, J.D. (2018). Behavioral Phenotyping of an Improved Mouse Model of Phelan-McDermid Syndrome with a Complete Deletion of the Shank3 Gene. eNeuro 5. 10.1523/ENEURO.0046-18.2018.

34. Levy, D.R., Tamir, T., Kaufman, M., Parabucki, A., Weissbrod, A., Schneidman, E., and Yizhar, O. (2019). Dynamics of social representation in the mouse prefrontal cortex. Nat Neurosci 22, 2013–2022. 10.1038/s41593-019-0531-z.

35. Sherwin, E., Bordenstein, S.R., Quinn, J.L., Dinan, T.G., and Cryan, J.F. (2019). Microbiota and the social brain. Science 366. 10.1126/science.aar2016.

36. Fortier, A.V., Meisner, O.C., Nair, A.R., and Chang, S.W.C. (2022). Prefrontal circuits guiding social preference: Implications in autism spectrum disorder. Neurosci Biobehav Rev 141, 104803. 10.1016/j.neubiorev.2022.104803.

37. Pagani, M., Bertero, A., Liska, A., Galbusera, A., Sabbioni, M., Barsotti, N., Colenbier, N., Marinazzo, D., Scattoni, M.L., Pasqualetti, M., et al. (2019). Deletion of Autism Risk Gene Shank3 Disrupts Prefrontal Connectivity. J Neurosci 39, 5299–5310. 10.1523/JNEUROSCI.2529-18.2019.

38. Yoo, T., Yoo, Y.-E., Kang, H., and Kim, E. (2022). Age, brain region, and gene dosage-differential transcriptomic changes in Shank3 -mutant mice. Front Mol Neurosci 15, 1017512. 10.3389/fnmol.2022.1017512.

39. Jacot-Descombes, S., Keshav, N.U., Dickstein, D.L., Wicinski, B., Janssen, W.G.M., Hiester, L.L., Sarfo, E.K., Warda, T., Fam, M.M., Harony-Nicolas, H., et al. (2020). Altered synaptic ultrastructure in the prefrontal cortex of Shank3 -deficient rats. Mol Autism 11, 89. 10.1186/s13229-020-00393-8.

40. Gacias, M., Gaspari, S., Santos, P.-M.G., Tamburini, S., Andrade, M., Zhang, F., Shen, N., Tolstikov, V., Kiebish, M.A., Dupree, J.L., et al. (2016). Microbiota-driven transcriptional changes in prefrontal cortex override genetic differences in social behavior. Elife 5. 10.7554/eLife.13442.

41. Hoban, A.E., Stilling, R.M., Ryan, F.J., Shanahan, F., Dinan, T.G., Claesson, M.J., Clarke, G., and Cryan, J.F. (2016). Regulation of prefrontal cortex myelination by the microbiota. Transl Psychiatry 6, e774. 10.1038/tp.2016.42.

42. Soliman, M.L., and Rosenberger, T.A. (2011). Acetate supplementation increases brain histone acetylation and inhibits histone deacetylase activity and expression. Mol Cell Biochem 352, 173–180. 10.1007/s11010-011-0751-3.

43. Mews, P., Egervari, G., Nativio, R., Sidoli, S., Donahue, G., Lombroso, S.I., Alexander, D.C., Riesche, S.L., Heller, E.A., Nestler, E.J., et al. (2019). Alcohol metabolism contributes to brain histone acetylation. Nature 574, 717–721. 10.1038/s41586-019-1700-7.

44. Frost, G., Sleeth, M.L., Sahuri-Arisoylu, M., Lizarbe, B., Cerdan, S., Brody, L., Anastasovska, J., Ghourab, S., Hankir, M., Zhang, S., et al. (2014). The short-chain fatty acid acetate reduces appetite via a central homeostatic mechanism. Nat Commun 5, 3611. 10.1038/ncomms4611.

45. Langfelder, P., and Horvath, S. (2008). WGCNA: an R package for weighted correlation network analysis. BMC Bioinformatics 9, 559. 10.1186/1471-2105-9-559.

46. Vuong, H.E., and Hsiao, E.Y. (2017). Emerging Roles for the Gut Microbiome in Autism Spectrum Disorder. Biol. Psychiatry 81, 411–423. 10.1016/j.biopsych.2016.08.024.

47. Iannone, L.F., Preda, A., Blottière, H.M., Clarke, G., Albani, D., Belcastro, V., Carotenuto, M., Cattaneo, A., Citraro, R., Ferraris, C., et al. (2019). Microbiota-gut brain axis involvement in neuropsychiatric disorders. Expert Rev Neurother 19, 1037–1050. 10.1080/14737175.2019.1638763.

48. Partrick, K.A., Chassaing, B., Beach, L.Q., McCann, K.E., Gewirtz, A.T., and Huhman, K.L. (2018). Acute and repeated exposure to social stress reduces gut microbiota diversity in Syrian hamsters. Behavioural Brain Research 345, 39–48. 10.1016/j.bbr.2018.02.005.

49. Machiels, K., Joossens, M., Sabino, J., De Preter, V., Arijs, I., Eeckhaut, V., Ballet, V., Claes, K., Van Immerseel, F., Verbeke, K., et al. (2014). A decrease of the butyrate-producing species Roseburia hominis and Faecalibacterium prausnitzii defines dysbiosis in patients with ulcerative colitis. Gut 63, 1275–1283. 10.1136/gutjnl-2013-304833.

50. Sauer, A.K., Bockmann, J., Steinestel, K., Boeckers, T.M., and Grabrucker, A.M. (2019). Altered Intestinal Morphology and Microbiota Composition in the Autism Spectrum Disorders Associated SHANK3 Mouse Model. International Journal of Molecular Sciences 20, 2134. 10.3390/ijms20092134.

51. De Angelis, M., Piccolo, M., Vannini, L., Siragusa, S., De Giacomo, A., Serrazzanetti, D.I., Cristofori, F., Guerzoni, M.E., Gobbetti, M., and Francavilla, R. (2013). Fecal microbiota and metabolome of children with autism and pervasive developmental disorder not otherwise specified. PLoS ONE 8, e76993. 10.1371/journal.pone.0076993.

52. Zhang, M., Ma, W., Zhang, J., He, Y., and Wang, J. (2018). Analysis of gut microbiota profiles and microbe-disease associations in children with autism spectrum disorders in China. Sci Rep 8, 13981. 10.1038/s41598-018-32219-2.

53. Strati, F., Cavalieri, D., Albanese, D., De Felice, C., Donati, C., Hayek, J., Jousson, O., Leoncini, S., Renzi, D., Calabrò, A., et al. (2017). New evidences on the altered gut microbiota in autism spectrum disorders. Microbiome 5, 24. 10.1186/s40168-017-0242-1.

54. Tomova, A., Husarova, V., Lakatosova, S., Bakos, J., Vlkova, B., Babinska, K., and Ostatnikova, D. (2015). Gastrointestinal microbiota in children with autism in Slovakia. Physiology & Behavior 138, 179–187. 10.1016/j.physbeh.2014.10.033.

55. James, D.M., Kozol, R.A., Kajiwara, Y., Wahl, A.L., Storrs, E.C., Buxbaum, J.D., Klein, M., Moshiree, B., and Dallman, J.E. (2019). Intestinal dysmotility in a zebrafish (Danio rerio) Shank3 a; Shank3 b mutant model of autism. Mol Autism 10, 3. 10.1186/s13229-018-0250-4.

56. Zierer, J., Jackson, M.A., Kastenmüller, G., Mangino, M., Long, T., Telenti, A., Mohney, R.P., Small, K.S., Bell, J.T., Steves, C.J., et al. (2018). The fecal metabolome as a functional readout of the gut microbiome. Nat. Genet. 50, 790–795. 10.1038/s41588-018-0135-7.

57. Schroeder, B.O., and Bäckhed, F. (2016). Signals from the gut microbiota to distant organs in physiology and disease. Nat. Med. 22, 1079–1089. 10.1038/nm.4185.

58. Braniste, V., Al-Asmakh, M., Kowal, C., Anuar, F., Abbaspour, A., Tóth, M., Korecka, A., Bakocevic, N., Ng, L.G., Guan, N.L., et al. (2014). The gut microbiota influences blood-brain barrier permeability in mice. Sci Transl Med 6, 263ra158. 10.1126/scitranslmed.3009759.

59. Khemir, S., Halayem, S., Azzouz, H., Siala, H., Ferchichi, M., Guedria, A., Bedoui, A., Abdelhak, S., Messaoud, T., Tebib, N., et al. (2016). Autism in Phenylketonuria Patients: From Clinical Presentation to Molecular Defects. J. Child Neurol. 31, 843–849. 10.1177/0883073815623636.

60. Gevi, F., Zolla, L., Gabriele, S., and Persico, A.M. (2016). Urinary metabolomics of young Italian autistic children supports abnormal tryptophan and purine metabolism. Mol Autism 7, 47. 10.1186/s13229-016-0109-5.

61. Alexander, D.C., Corman, T., Mendoza, M., Glass, A., Belity, T., Wu, R., Campbell, R.R., Han, J., Keiser, A.A., Winkler, J., et al. (2022). Targeting acetyl-CoA metabolism attenuates the formation of fear memories through reduced activity-dependent histone acetylation. Proceedings of the National Academy of Sciences 119, e2114758119. 10.1073/pnas.2114758119.

62. Mews, P., Donahue, G., Drake, A.M., Luczak, V., Abel, T., and Berger, S.L. (2017). Acetyl-CoA synthetase regulates histone acetylation and hippocampal memory. Nature 546, 381–386. 10.1038/nature22405.

63. Hofford, R.S., Mervosh, N.L., Euston, T.J., Meckel, K.R., Orr, A.T., and Kiraly, D.D. (2021). Alterations in microbiome composition and metabolic byproducts drive behavioral and transcriptional responses to morphine. Neuropsychopharmacology. 10.1038/s41386-021-01043-0.

64. Caporaso, J.G., Kuczynski, J., Stombaugh, J., Bittinger, K., Bushman, F.D., Costello, E.K., Fierer, N., Peña, A.G., Goodrich, J.K., Gordon, J.I., et al. (2010). QIIME allows analysis of high-throughput community sequencing data. Nat. Methods 7, 335–336. 10.1038/nmeth.f.303.

65. Clarke, E.L., Taylor, L.J., Zhao, C., Connell, A., Lee, J.-J., Fett, B., Bushman, F.D., and Bittinger, K. (2019). Sunbeam: an extensible pipeline for analyzing metagenomic sequencing experiments. Microbiome 7, 46. 10.1186/s40168-019-0658-x.

66. Bolger, A.M., Lohse, M., and Usadel, B. (2014). Trimmomatic: a flexible trimmer for Illumina sequence data. Bioinformatics 30, 2114–2120. 10.1093/bioinformatics/btu170.

67. Li, H., and Durbin, R. (2009). Fast and accurate short read alignment with Burrows-Wheeler transform. Bioinformatics 25, 1754–1760. 10.1093/bioinformatics/btp324.

68. Buchfink, B., Xie, C., and Huson, D.H. (2015). Fast and sensitive protein alignment using DIAMOND. Nat Methods 12, 59–60. 10.1038/nmeth.3176.

69. Kanehisa, M., and Goto, S. (2000). KEGG: kyoto encyclopedia of genes and genomes. Nucleic Acids Res 28, 27–30. 10.1093/nar/28.1.27.

70. Love, M.I., Huber, W., and Anders, S. (2014). Moderated estimation of fold change and dispersion for RNA-seq data with DESeq2. Genome Biol 15, 550. 10.1186/s13059-014-0550-8.

71. Silverman, J.L., Yang, M., Lord, C., and Crawley, J.N. (2010). Behavioural phenotyping assays for mouse models of autism. Nat. Rev. Neurosci. 11, 490–502. 10.1038/nrn2851.

72. Torre, D., Lachmann, A., and Ma’ayan, A. (2018). BioJupies: Automated Generation of Interactive Notebooks for RNA-Seq Data Analysis in the Cloud. cels 7, 556–561.e3. 10.1016/j.cels.2018.10.007.

73. Ritchie, M.E., Phipson, B., Wu, D., Hu, Y., Law, C.W., Shi, W., and Smyth, G.K. (2015). limma powers differential expression analyses for RNA-sequencing and microarray studies. Nucleic Acids Res. 43, e47. 10.1093/nar/gkv007.

74. Raudvere, U., Kolberg, L., Kuzmin, I., Arak, T., Adler, P., Peterson, H., and Vilo, J. (2019). g:Profiler: a web server for functional enrichment analysis and conversions of gene lists (2019 update). Nucleic Acids Res. 47, W191–W198. 10.1093/nar/gkz369.

75. Krämer, A., Green, J., Pollard, J., and Tugendreich, S. (2014). Causal analysis approaches in Ingenuity Pathway Analysis. Bioinformatics 30, 523–530. 10.1093/bioinformatics/btt703.

76. Zhao, S., Li, H., Han, W., Chan, W., and Li, L. (2019). Metabolomic Coverage of Chemical-Group-Submetabolome Analysis: Group Classification and Four-Channel Chemical Isotope Labeling LC-MS. Anal. Chem. 91, 12108–12115. 10.1021/acs.analchem.9b03431.

77. Wu, Y., and Li, L. (2016). Sample normalization methods in quantitative metabolomics. Journal of Chromatography A 1430, 80–95. 10.1016/j.chroma.2015.12.007.

78. Zhou, R., Tseng, C.-L., Huan, T., and Li, L. (2014). IsoMS: Automated Processing of LC-MS Data Generated by a Chemical Isotope Labeling Metabolomics Platform. Anal. Chem. 86, 4675–4679. 10.1021/ac5009089.

79. Li, L., Li, R., Zhou, J., Zuniga, A., Stanislaus, A.E., Wu, Y., Huan, T., Zheng, J., Shi, Y., Wishart, D.S., et al. (2013). MyCompoundID: Using an Evidence-Based Metabolome Library for Metabolite Identification. Anal. Chem. 85, 3401–3408. 10.1021/ac400099b.

80. Pang, Z., Chong, J., Zhou, G., de Lima Morais, D.A., Chang, L., Barrette, M., Gauthier, C., Jacques, P.-É., Li, S., and Xia, J. (2021). MetaboAnalyst 5.0: narrowing the gap between raw spectra and functional insights. Nucleic Acids Research 49, W388–W396. 10.1093/nar/gkab382.

81. Breen, M.S., Fan, X., Levy, T., Pollak, R.M., Collins, B., Osman, A., Tocheva, A.S., Sahin, M., Berry-Kravis, E., Soorya, L., et al. (2023). Large 22q13.3 deletions perturb peripheral transcriptomic and metabolomic profiles in Phelan-McDermid syndrome. Human Genetics and Genomics Advances 4, 100145. 10.1016/j.xhgg.2022.100145.

82. Dehaven, C.D., Evans, A.M., Dai, H., and Lawton, K.A. (2010). Organization of GC/MS and LC/MS metabolomics data into chemical libraries. J Cheminform 2, 9. 10.1186/1758-2946-2-9.

83. Evans, A.M., DeHaven, C.D., Barrett, T., Mitchell, M., and Milgram, E. (2009). Integrated, nontargeted ultrahigh performance liquid chromatography/electrospray ionization tandem mass spectrometry platform for the identification and relative quantification of the small-molecule complement of biological systems. Anal Chem 81, 6656–6667. 10.1021/ac901536h.

