## Supplemental Figures for "Acetate supplementation rescues social deficits and alters transcriptional regulation in prefrontal cortex of Shank3 deficient mice"

### Supplementary Figures

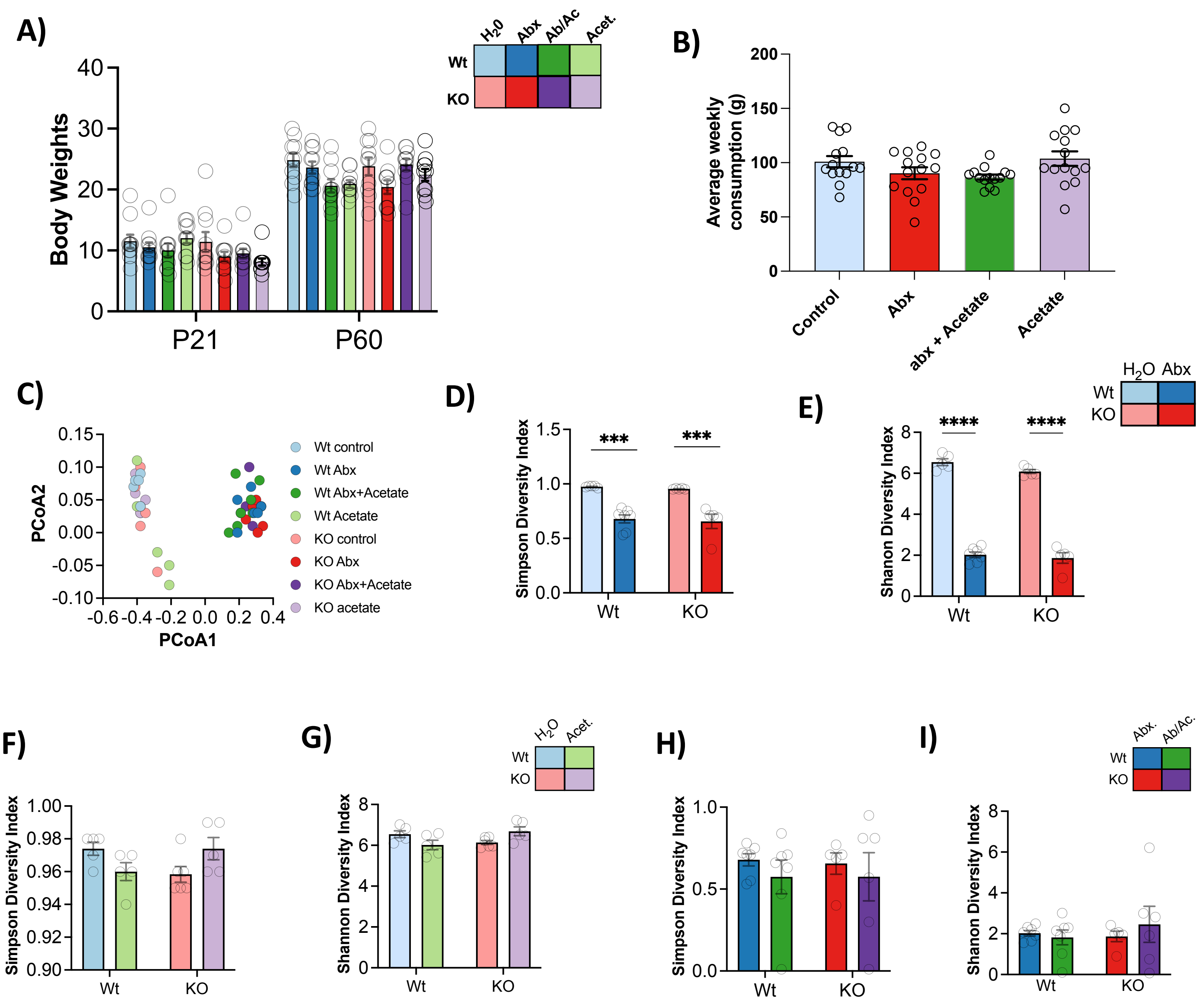


**Supp Figure 1: Acetate Replenishment Does not Affect Overall Animal Health (A)** Wild-type and Shank3$^{KO}$ mice show normal weight gain from PND21 to PND60 regardless of genotype or treatment **(B)** Animals of all genotypes drink control water, antibiotic treated (Abx) antibiotic plus acetate (Abx+ acetate) and acetate treated water at similar rates over development. All data shown as SEM **(C)** Unweighted PCOA plot of beta diversity shows a marked shift induced by antibiotic treatment in both genotypes **(D)** and **(E)** Antibiotics results in marked depletion of microbial diversity in both Wt and Shank3$^{KO}$ mice **(F)** and **(G)** Acetate treatment interactions with genotype to alter microbiome diversity **(H)** and **(I)** Treatment with acetate does not reverse Abx effects on microbiome diversity

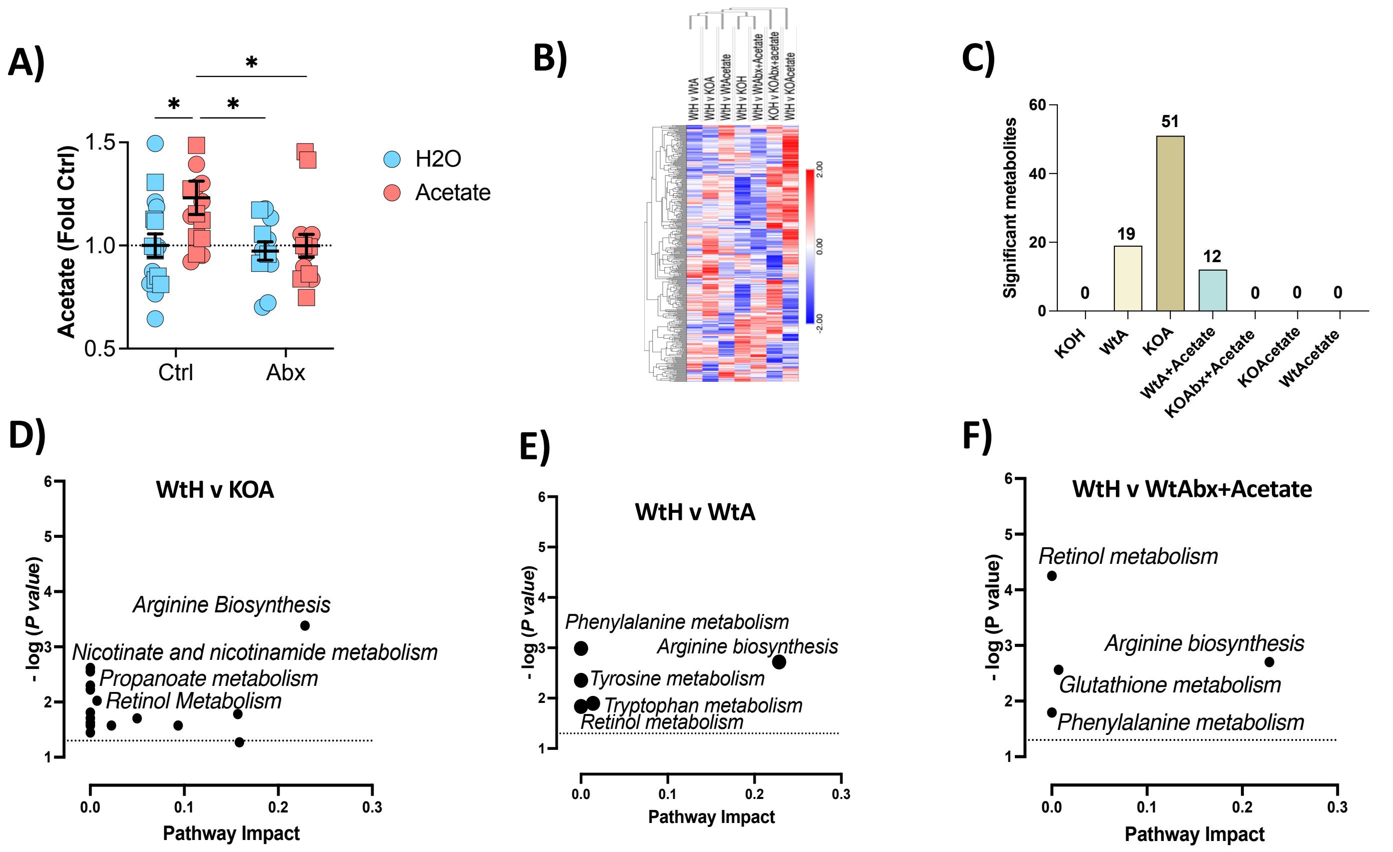


**Supp Figure 2: Acetate Replenishment Alters Gut Microbial Diversity and Normalizes Abx Effects on Serum Metabolic Profile (A)** Targeted SCFA metabolomic analysis reveals main effect of acetate treatment in both Wt and KO animals compared to control treated counterpart. Abx treatment was found to decrease levels of acetate in both genotypes, even in animals treated with Abx +acetate [Two way ANOVA Holm-Sidak's post-hoc] **(B)** Unsupervised hierarchal clustering and heatmap (Blue, low and red ,high) depiction of relative abundance of the 372 metabolites identified across all treatment groups **(C)** The total number of significant differentially expressed metabolites ( y axis) for each treatment group (x axis) ($p$<0.05 q<0.25) **(D)** Pathway analysis of differentially expressed metabolites in KO Abx treated animals (KOA) compared to WtH reveals significant pathway enrichment (y axis; –log10 p value) for Nicotinate and nicotinamide metabolism and 19 other metabolism pathways relative to pathway impact ( x axis) **(E)** Pathway analysis of differentially expressed metabolites in Wt Abx treated animals (WtA) compared to Wt controls (WtH) reveals significant pathway enrichment (y axis; –log10 p value) for Phenylalanine metabolism and four other metabolism pathways relative to pathway impact ( x axis) **(F)** Pathway analysis of differentially expressed metabolites in Wt Abx + acetate treated animals (WtAbx+ Acetate) compared to WtH reveals significant pathway enrichment (y axis; –log10 p value) for Retinol metabolism and three other metabolism pathways relative to pathway impact ( x axis). Pathway impact is a combination of the centrality and pathway enrichment results computed by adding the importance measures of each of matched metabolite and dividing by the sum of the importance measures of all metabolites in each pathway. All data shown as SEM. *$p$ < 0.05; **$p$ < 0.01; ***$p$ < 0.001; ****$p$ < 0.0001]

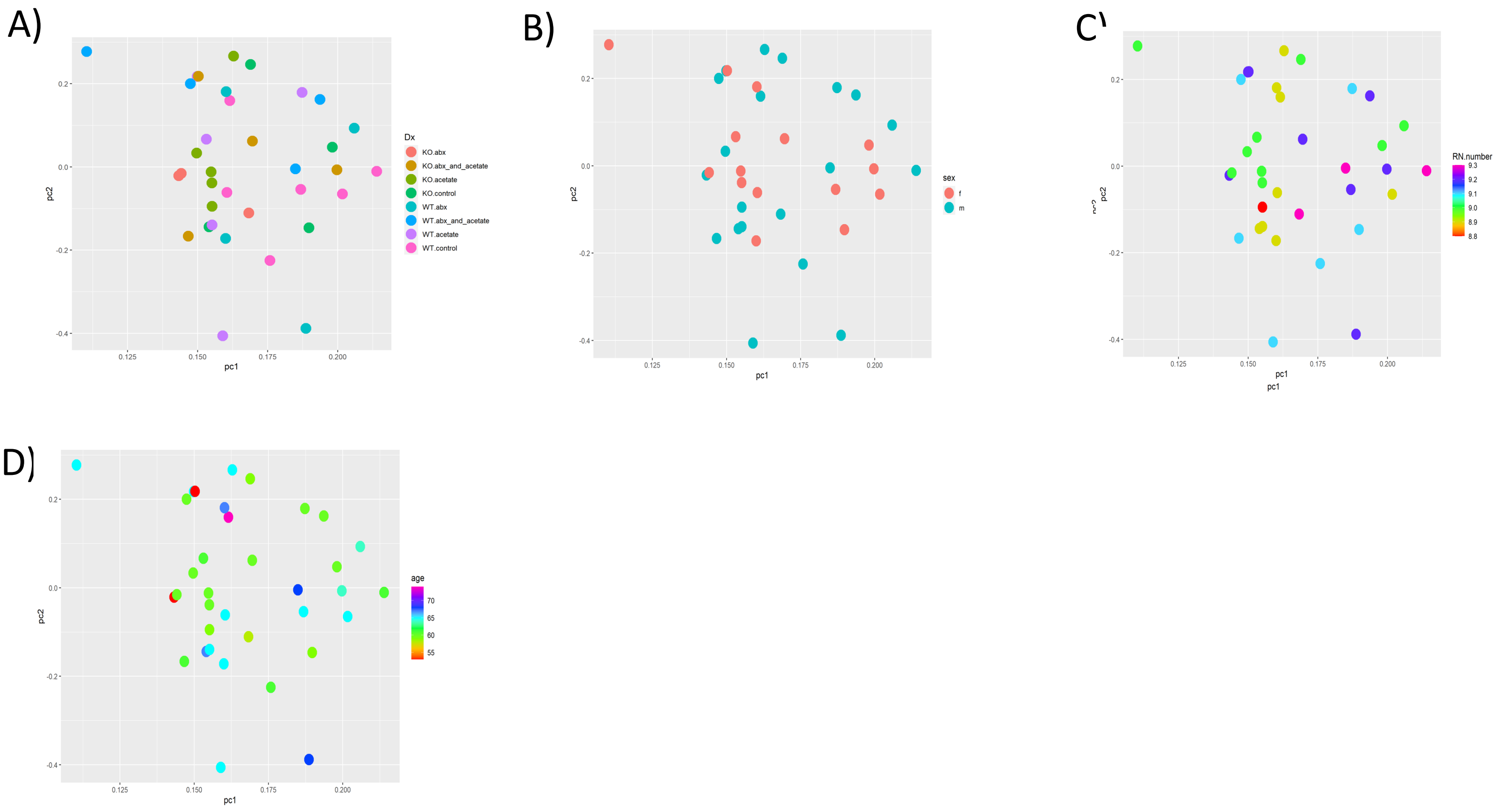


**Supp Figure 3:Supplemental RNA Sequencing Figures (A)** Principle Component Analysis (PCA) does not show clustering by genotype or treatment **(B)** Sex **(C)** RIN value or **(D)** age of animals

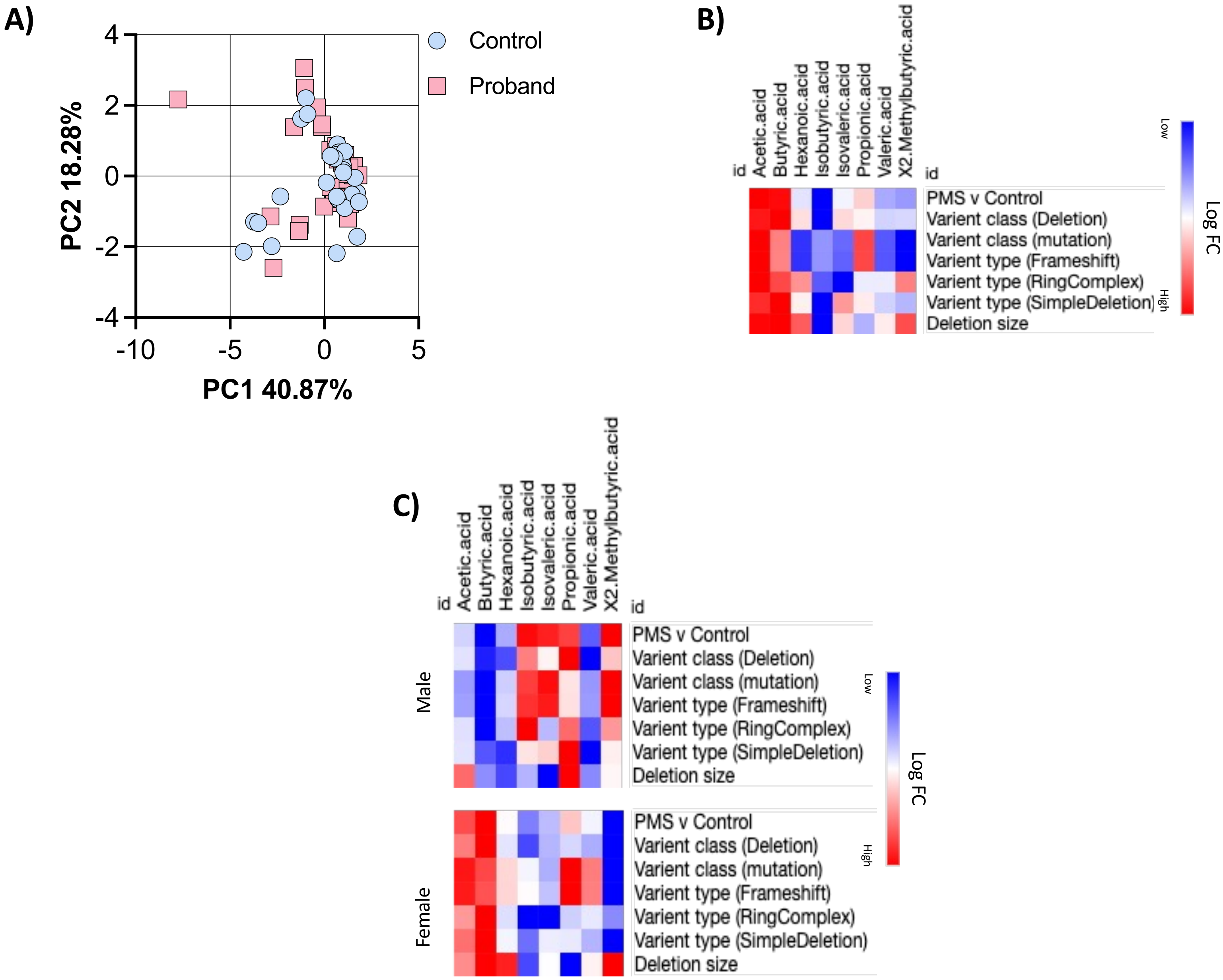


**Supp Figure 4: Levels of Serum SCFA are not Significantly Altered by PMS Genotype. (A)** PCA showing no clear separation between PMS and controls in levels of SCFAs **(B)** Heatmap showing correlations between levels of SCFA's and Genotype Varient class, Varient type and Deletion size in both males and female PMS v control counterparts **(C)** Heatmap showing correlations between levels of SCFA's and Genotype Varient class, Varient type and Deletion size in males and female PMS patients separately.

| Table S11. Clinical features of Participants in Targeted Metabolomics Study | |
|---|---|
| Demographics for both Control and Proband | Percentage (%) or Mean ± SD |
| Age months | 139.7 ± 105.00 |
| Sex, Male (N,%) | 43.33% |
| Ethnicity, white (N, %) | 78.33% |
| Clinical Indices (Probands only) | Percentage (%) or Mean ± SD |
| Meds.present (n=32) % | 78.13% |
| ADOS.meet.criteria (n=31) % | 80.65% |
| ADI.meet.criteria (n=29) % | 62.07% |
| GI.present (n=26) % | 69.23% |
| Number.meds (n=32) | 4.28 ± 4.73 |
| Verbal.IQ.DQ (n=30) | 25.08 ± 19.91 |
| nonverbal.IQ.DQ (n=29) | 29.62 ± 18.04 |
| Full.scale.IQ.DQ (n=29) | 26.71 ± 18.33 |
| ADOS.total (n=30) | 16.27 ± 6.90 |
| ADOS.SA (n=30) | 12.53 ± 5.65 |
| ADOS.RRB (n=30) | 3.73 ± 2.21 |
| ADOS.severity (n=29) | 6.38 ± 2.27 |
| ADI.social (n=26) | 17.92 ± 8.32 |
| ADI.comm (n=27) | 11.35 ± 4.11 |
| ADI.RRB (n=27) | 3.69 ± 1.91 |
| VABS.comm (n=30) | 50.20 ± 16.03 |
| VABS.DLS (n=30) | 54.93± 15.73 |
| VABS.social (n=30) | 59.50 ± 16.03 |
| VABS.motor (n=28) | 58.33 ± 14.54 |
| VABS.adapt.comp (n=30) | 53.70 ± 14.44 |
| ABC.irritability (n=29) | 8.34 ± 7.34 |
| ABC.lethargy (n=29) | 9.38 ± 7.75 |
| ABC.stereotypy (n=29) | 4.66 ± 4.75 |
| ABC.hyperactivity (n=29) | 20.48 ± 12.81 |
| ABC.ppropriate.speech (n=28) | 2.21 ± 2.87 |
| Abbreviations: Aberrant Behavior Checklist (ABC); Autism Diagnostic Interview (ADI); Autism Diagnostic Observation Schedule (ADOS); Developmental Quotient (DQ); Intellectual Quotient (IQ); Restrictive and Repetitive Behavior (RRB); Social Affect (SA); Vineland Adaptive Behavior Scales (VABS).GI; Gastrointestinal tract | |
